# Temperature-dependent regulation of bacterial cell division hydrolases by the coordinated action of a regulatory RNA and the ClpXP protease

**DOI:** 10.1101/2025.04.01.646552

**Authors:** Viktor H. Mebus, Larissa M. Busch, Morten Børre, Tobias K. Nielsen, Martin Saxtorph Bojer, Camilla Henriksen, Maria D. Barbuti, Danae M. Angeles, Kamilla Brejndal, Stephan Michalik, Manuela Gesell Salazar, Morten Kjos, Uwe Völker, Birgitte H. Kallipolitis, Dorte Frees

**Affiliations:** Department of Veterinary and Animal Sciences, Faculty of Health and Medical Sciences, University of Copenhagen, Copenhagen, Denmark; Department of Functional Genomics, Interfaculty Institute for Genetics and Functional Genomics, University Medicine Greifswald, Greifswald, Germany; Department of Biochemistry and Molecular Biology, Faculty of Natural Sciences, University of Southern Denmark, Odense, Denmark; Faculty of Chemistry, Biotechnology and Food Sciences, Norwegian University of Life Sciences, Norway

## Abstract

A defining feature of bacteria is the peptidoglycan cell wall which provides structural integrity and prevents osmotic lysis. While peptidoglycan hydrolases are required for daughter cell separation, dysregulated cell wall degradation may result in cell lysis. The mechanisms allowing bacteria to control these deadly enzymes in response to environmental changes remain incompletely understood. Here, we find that in *Staphylococcus aureus*, temperature-dependent regulation of such hydrolases occurs by the coordinated action of a CHAP domain-specific regulatory RNA and the ClpXP protease. Using a proteomics approach, we identify a hitherto uncharacterized <u>C</u>lp<u>X</u>P <u>c</u>ontrolled <u>a</u>utolysin, CxcA, with a catalytic CHAP domain and show that it contributes to separation of daughter cells. CxcA is positively controlled by a non-coding RNA, named Rbc1 (for RNA <u>b</u>inding to CHAP domain) transcribed from the antisense strand of *cxcA*. Notably, Rbc1 is capable of base pairing with RNAs encoding the CHAP domains of numerous cell wall hydrolases and we show that Rbc1 works *in trans* to upregulate the cell division hydrolase Sle1. Specifically, Rbc1 functions as a thermosensor allowing for upregulation of CxcA and Sle1 at low temperature where daughter cell separation is impeded. Interestingly, the Rbc1-mediated up-regulation of CxcA and Sle1 does not involve mRNA stabilization or increased translation; instead, Rbc1 depletion increases ClpXP-mediated degradation. In conclusion, we identify a novel cell division hydrolase that is highly conserved in Staphylococci and show that it is co-regulated with enzymes containing the catalytic CHAP domain via transcriptional regulation, an RNA-RNA temperature sensory mechanism and the ClpXP protease.

## Introduction

The cell wall is essential for bacterial survival and it is the target of β-lactams, the most prescribed antibiotics for treatment of bacterial infections (ECDC, 2019; Schneider & Sahl, 2010). This important class of antibiotics includes the penicillins, cephalosporins, and carbapenems that all possess a reactive four-membered ring that interfere with synthesis of the bacterial cell wall by binding and inhibiting the activity of a group of enzymes called penicillin-binding proteins (PBPs) (Tipper & Strominger, 1965). Bacterial cell death following β -lactam treatment has been linked to the continued activity of cell wall degrading enzymes (Cho *et al*., 2014; Kitano & Tomasz, 1979; Salamaga *et al*., 2021). While cell wall hydrolases play pivotal roles in facilitating the integration of new cell wall material and in daughter cell separation, dysregulated degradation of the cell wall may lead to catastrophic cell lysis. Hence, bacteria use sophisticated control mechanisms to control abundance and activity of these potentially deadly enzymes in space and time (Do *et al*., 2020; Vollmer *et al*., 2008). The regulatory mechanisms governing the protein levels of autolysins in response to changes in growth conditions, however, remain far less explored.

In nearly all bacteria, the cell wall consists of a peptidoglycan (PG) meshwork composed of linear glycan strands cross-linked by short peptide chains (Silhavy *et al*., 2010). Gram-negative bacteria have a thin PG layer located between the inner and outer membrane, whereas the cell wall of Gram-positive bacteria is composed of a single thick layer of PG decorated with teichoic acids (TAs) that are unique to Gram-positive cells (Silhavy *et al*., 2010). The synthesis of PG is facilitated by two functionally distinct protein complexes: the divisome which constructs the septal cross wall during cell division, and the elongasome, which enables bacterial elongation by synthesizing PG along the length of the cell (Rohs & Bernhardt, 2021). In both places, the synthesis of PG is coordinated with cell wall hydrolase activity. These hydrolases, classified by their enzymatic activity, include glycosidases (cleavage within the glycan strands), peptidases (cleavage within the stem peptide or the cross-bridge), and amidases (cleave the bond between the stem peptide and the glycan strand) (Do *et al*., 2020; Vollmer *et al*., 2008). PG hydrolases typically display a modular architecture where highly conserved catalytic domains encoding the different enzymatic activities are combined with different cell wall-binding domains (Vermassen *et al*., 2019a).

The opportunistic pathogen *Staphylococcus aureus* colonizes the anterior nares and the skin of approximately 30% of the human population (Krismer *et al*., 2017; Liu *et al*., 2015). Colonization is most often asymptomatic, however, self-inoculation can give rise to fatal infections of the deeper tissues and the blood (Tong *et al*., 2015). Since the temperature of the skin and nasal cavity is estimated to be lower than the body temperature, respectively, 32°C and 34°C, the transition from colonization to infection is associated with an upshift in temperature (Bastock *et al*., 2021; Costa *et al*., 2024; Keck *et al*., 2000). In recent years, *S. aureus* has become a model organism for studying cell division in coccoid bacteria and despite its coccoid shape, *S. aureus* similar to rod-shaped bacteria possesses two distinct machineries for cell division and cell wall elongation (Pinho & Foster, 2024; Reichmann *et al*., 2019). The *S. aureus* core genome is predicted to encode at least eighteen PG degrading enzymes and about half of these enzymes remain functionally uncharacterized (Wang *et al*., 2022; Wang *et al*., 2024). Most of the *S. aureus* PG hydrolases contain the highly conserved C-terminal **c**ysteine, **h**istidine-dependent **a**midohydrolase**/p**eptidase (CHAP) domain that has peptidase and/or amidase activity (Bateman & Rawlings, 2003; Mitchell *et al*., 2021; Wang *et al*., 2022; Wang *et al*., 2024). One well-characterized example is the Sle1 amidase that is associated with separation of *S. aureus* daughter cells during cell division (Monteiro *et al*., 2015; Zhou *et al*., 2015; Thalsø-Madsen *et al*., 2019). According to the current paradigm, *S. aureus* builds a septal cross-wall, generating two hemispherical daughter cells that separate by ultrafast popping when the peripheral septal ring is resolved in a process involving Sle1 and mechanical crack propagation (Monteiro *et al*., 2015; Zhou *et al*., 2015; Thalsø-Madsen *et al*., 2019). The Sle1 cell wall hydrolase is a substrate of the highly conserved cytoplasmic protease, ClpXP that in *S. aureus* performs regulated proteolysis of proteins that need to be controlled in space and time (Feng *et al*., 2013; Stahlhut *et al*., 2017). In *S. aureus* cells devoid of ClpXP activity, the accumulation of Sle1 elicits splitting of premature daughter cells leading to fatal cell lysis, hence, underscoring the critical importance of regulating autolytic activity (Henriksen *et al*., 2024; Jensen *et al*., 2019).

In the current study, we identify the uncharacterized cell wall hydrolase SAOUHSC_00671 (SAUSA300_0651) as a novel substrate of the ClpXP protease and accordingly named it CxcA for <u>C</u>lp<u>X</u>P <u>c</u>ontrolled <u>a</u>utolysin. CxcA has the same modular organization as Sle1 including a C-terminal CHAP domain, and we show that Sle1 and CxcA are co-regulated by Rbc1, a non-coding RNA transcribed from the anti-sense strand of the *cxcA* gene. *In vitro*, Rbc1 is capable of binding to the CHAP encoding region of at least three more transcripts encoding cell wall hydrolases (SsaA, EssH, and SAUSA300_02503) indicating that *S. aureus* may employ the same anti-sense RNAs to coordinate a class of enzymes containing the same catalytic domain. Specifically, Rbc1 functions as a thermosensor allowing *S. aureus* to enrich cell division hydrolases in response to decreasing temperature where separation of *S. aureus* daughter cells becomes delayed relative to septum synthesis. In summary, we demonstrate that CxcA has autolytic activity and that its abundance is tightly regulated at the level of transcription, translation, and post-translationally.

## Results

### Proteomics reveals ClpXP-mediated degradation of the uncharacterized autolysin

SAOUHSC_00671/ SAUSA300_0651

In *S. aureus*, the ClpXP protease performs regulated proteolysis of proteins that, like Sle1, need to be controlled in space and time (Stahlhut *et al*., 2017). The cell division autolysin, Sle1 was first identified as a ClpP substrate when identifying proteins co-purifying with a His-tagged proteolytic inactive ClpP^TRAP^ (Feng *et al*., 2013). Additionally, three predicted autolysins were captured as putative ClpP substrates, indicating that more autolysins could be regulated by ClpP-proteolysis (Feng *et al*., 2013). To investigate if ClpXP-mediated degradation contributes to controlling more cell wall hydrolases than Sle1, we here used stable isotope labeling with amino acids in cell culture (SILAC) to identify proteins accumulating in *S. aureus* cells lacking ClpXP activity. The ClpXP activity was abolished by introducing an inactivating substitution in the ClpP recognition IGF motif of ClpX (ClpX_I265E_) thereby preventing formation of the ClpXP protease while preserving ClpX unfoldase activity (Stahlhut *et al*., 2017). Total cellular protein samples were extracted from cells harvested in exponential phase (OD_600_=0.4) (Fig. 1A) and in stationary phase (OD_600_=1.8) (Fig. S1) cultivated in pMEM. A total of 1453 proteins were identified, of which 1290, were identified with at least two peptides, which were then included into the subsequent search for proteins differentially abundant between *S. aureus* cells expressing the ClpX_I265E_ variant or the wild type ClpX variant, respectively (Supplemental Table S1). In total, 11 of the 18 predicted cell wall hydrolases of *S. aureus* (Wang *et al*., 2022) were identified with at least two peptides. Six predicted cell wall hydrolysases had significantly increased protein levels (fold change ≥ 2, q-value ≤ 0.05) in at least one growth phase in cells with inactivated ClpXP, namely, SsaA, Sle1, LytM, SceD, SAOUHSC_02576, and SAOUHSC_00671. For five of these proteins, the accumulation is associated with increased transcription in cells expressing the ClpX_I265E_ variant (Stahlhut *et al*., 2017). In contrast, no change in transcription was observed for the gene encoding SAOUHSC_00671, and as SAOUHSC_00671 was previously identified as a putative ClpP substrate (Feng *et al*., 2013), we decided to focus on this uncharacterized cell wall hydrolase that we renamed “CxcA” for <u>C</u>lp<u>X</u>P <u>c</u>ontrolled <u>a</u>utolysin.

**Figure 1.**
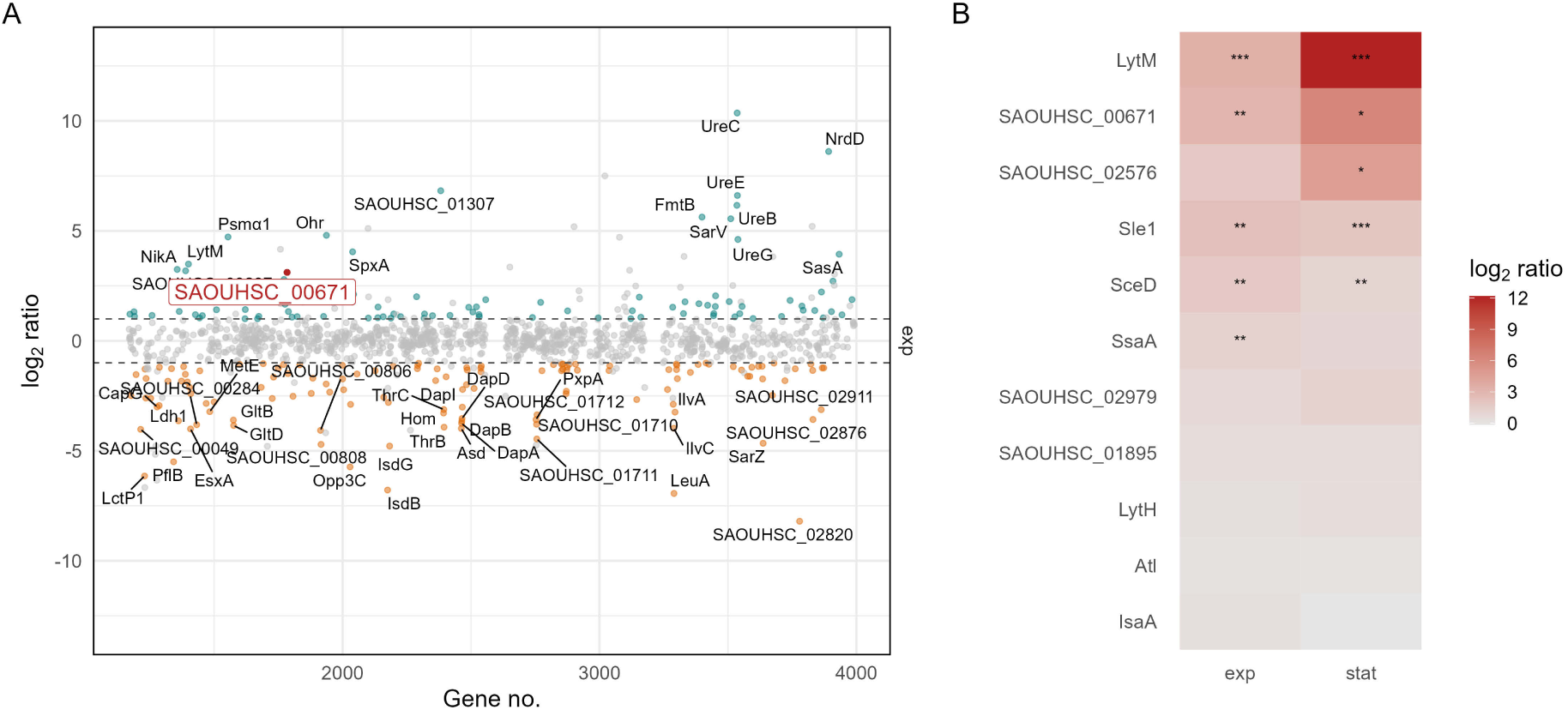
SAOUHSC_00671 autolysin is increased in protein abundance in S. aureus cells carrying the ClpX_I265E_ variant compared to cells carrying the wild-type ClpX variant. **(A)** Overview of the limmaDE statistics for the proteomic analysis of NCTC8325-4 ΔclpX cells complemented with the ClpX_I265E_ variant compared to cells complemented with the wild-type ClpX variant. Cells were harvested during exponential and stationary growth phase. Unique proteins are plotted as dots according to their genomic locus coordinate (grey: not significantly changed in protein abundance, cyan: significantly increased, orange: significantly decreased). Significantly changed in abundance is defined as the Benjamini-Hochberg adjusted P-value ≤ 0.05 and absolute log_2_ ratio ≥ 1. Significantly altered proteins with an absolute log_2_ ratio ≥ 3 are labeled. Autolysin SAOUHSC_00671 is highlighted in red. **(B)** Autolysins according to Wang et al. (2022) which are identified with at least two peptides are displayed. For significantly different protein abundances (adjusted P-value ≤ 0.05 and absolute log_2_ ratio ≥ 1), the adjusted p-value is indicated as following: *: ≤ 0.05, **: ≤ 0.01, ***: ≤ 0.001.

### Deletion of *cxcA* delays daughter cell splitting and mitigates autolysis

The *cxcA* gene is predicted to encode a 265 amino acid protein with a 25 amino acid N-terminal Sec-type signal peptide followed by two anchoring LysM domains (small globular protein domains associated with binding to the septal cross-wall) and a catalytic CHAP domain (Fig. 2A) (Bateman & Rawlings, 2003). The *cxcA* gene is known to be transcriptionally controlled by the essential two-component system, WalRK, that controls transcription of several *S. aureus* PG hydrolase genes, and to be upregulated by the <u>G</u>lycopeptide <u>Re</u>sistance <u>A</u>ssociated GraXRS-VraFG 5-component system that is localized downstream of the *cxcA* gene – see Fig. 2A (Delauné *et al*., 2012; Falord *et al*., 2011).

**Figure 2.**
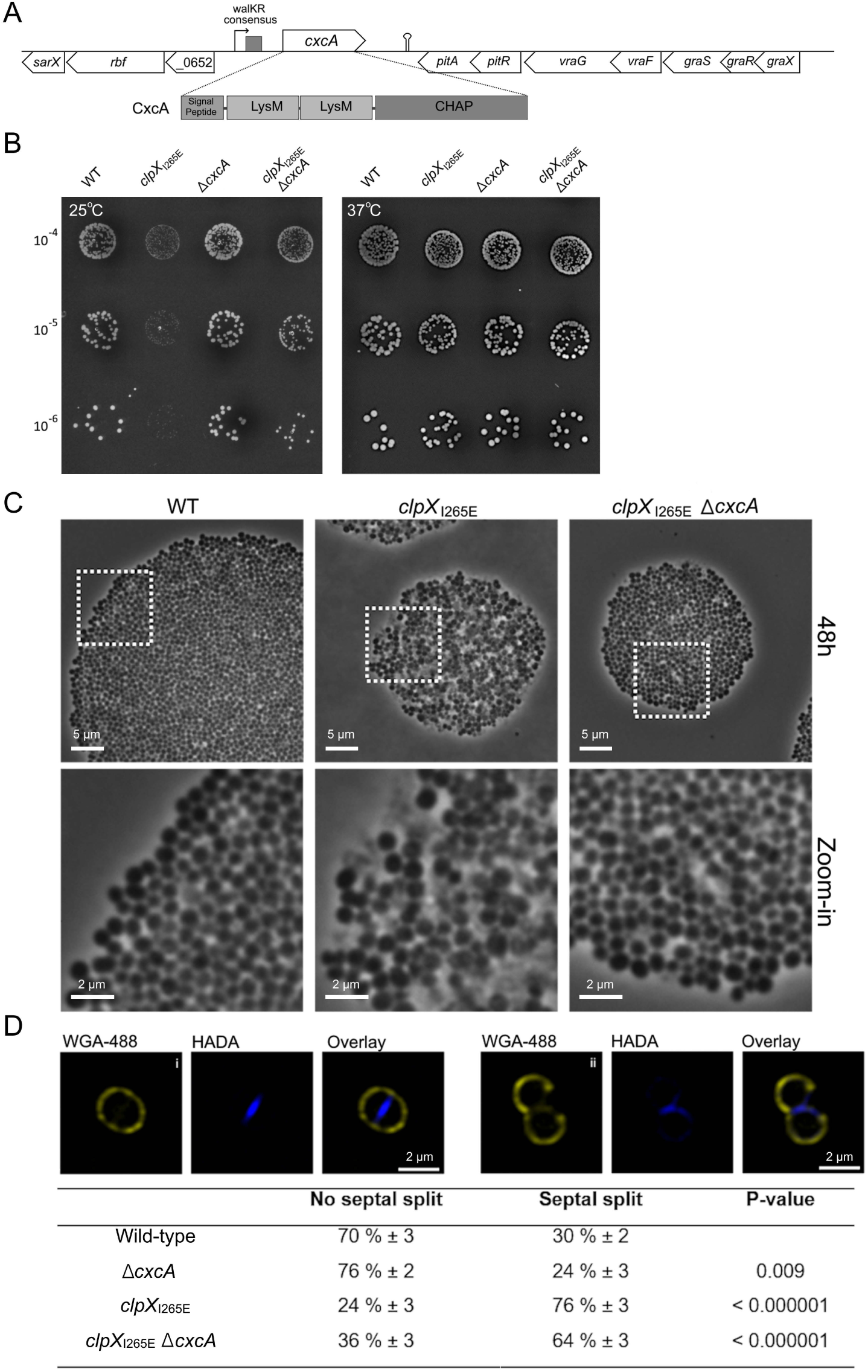
Inactivation of cxcA delays daughter cell separation and reduces lysis of cells devoid of ClpXP. **(A)** Schematic of the genomic and protein domain organization of CxcA. **(B)** Spot-dilution assay: WT and mutant strains were grown exponentially at 37°C in TSB. Subsequently, 10-fold serially diluted cells (OD_600_ ∼ 0.4) were spotted on TSA-agar plates that were incubated at 25°C or 37°C for 24 hours. **(C)** Growth of single colonies were studied by time-lapse microscopy at 25°C: Images were acquired every 5 minutes for at least 48 hours. **(D)** Daughter cell splitting was tracked by labeling with HADA (blue, label PG synthesis) and WGA-488 (yellow; binds to the outer cell wall but not the septum) before imaging with SR-SIM, a scale-bar of 2 µm is on the overlay image. For each strain, the proportion of cells that proceeded to daughter cell split, was quantified for at least 500 cells displaying a closed HADA-stained septum. Statistical analysis was performed using the chi square test for independence.

To investigate the function of this uncharacterized protein we used allelic exchange to create an in-frame deletion of the *cxcA* gene (preserving the nucleotides encoding the 12 first and the last 10 amino acids) in the preferred MRSA model strain, JE2 (Fey *et al*., 2013). In the JE2 WT background, deletion of the *cxcA* gene did neither affect growth rate in liquid media (the doubling times of the WT and Δ*cxcA* were 24.6 ± 1.4 min. and 24.6 ± 1.3 min respectively), nor did it reduce plating efficiency or colony size in spot-dilution assays (Fig. 2B). *S. aureus* cells lacking ClpXP activity grow normally at 37°C, but are unable to form colonies at 25°C (Jensen *et al*., 2019; Stahlhut *et al*., 2017). Interestingly, deletion of the *cxcA* gene enabled JE2 carrying the *clpX_I265E_* allele to form colonies at 25°C (Fig. 2B). When growth of single cells was followed by time lapse microscopy at 25°C, explosive lysis became highly prevalent in colonies formed by cells expressing the *clpX_I265E_* variant after 48h (Fig. 2C). Strikingly, lysis of JE2 *clpX_I265E_* cells was visibly rescued upon deletion of the *cxcA* gene (Fig. 2C), indicating that the CxcA autolysin contributes to the temperature dependent lysis observed for *S. aureus* cells lacking ClpXP activity.

To investigate if CxcA activity plays a role in autolytic splitting of WT daughter cells, we next employed super-resolution structured illumination microscopy (SR-SIM) on early exponential cells incubated for 10 min with the fluorescent wheat germ agglutinin (WGA-488) that specifically labels the peripheral cell wall (Monteiro *et al*., 2015). Additionally, cells were stained with the blue, fluorescent D-amino acid, hydroxycoumarin carbonyl amino-D-alanine (HADA), that labels region of active peptidoglycan synthesis (Kuru *et al*., 2012). This dual labeling allowed us to track daughter cell splitting in cells that had completed septum closure during the pulse-labeling with HADA (Fig. 2D). For each strain, we counted at least 500 cells with a closed HADA-stained septum and calculated the fraction displaying splitting of the HADA stained septum as illustrated in Fig. 2D. This analysis revealed that deletion of *cxcA* significantly reduced the fraction of daughter cells that separated in the time course of the experiment, both in the WT and the *clpX_I265E_* background (Fig. 2D). To further investigate the impact of CxcA on the *S. aureus* cell cycle, Nile Red stained cells imaged with SR-SIM were assigned to different phases of the cell cycle based on the state of septal ingrowths (Monteiro *et al*., 2015): newly separated daughter cells that have not initiated septum formation were assigned to phase 1, cells in the process of synthesizing division septa were assigned to phase 2, while cells displaying a closed septum were assigned to phase 3. This analysis revealed a slight, but significant increase in the fraction of cells displaying a closed septum upon deletion of the *cxcA* gene from 14.7 ± 2.0 % to 19.0 ± 1.0 %, (P < 0.005) in the JE2 WT background, and from 12.0 ± 0.7 % to 17.8 ± 0.7 % in JE2 cells lacking ClpXP activity (P < 10^-7^). Since the percentage of cells observed in each growth phase should be proportional to the time spent in this stage of the cell cycle, this result supports that CxcA promotes *S. aureus* daughter splitting.

### CxcA level is positively controlled by the Rbc1 anti-sense RNA

Previous transcriptome mapping revealed that a non-coding anti-sense (as) RNA is transcribed from the complementary strand of the *cxcA* gene, Fig. 3A (Beaume *et al*., 2010; Mäder *et al*., 2016). This asRNA was previously designated S254, or Teg20as, however, for reasons explained below, we renamed this asRNA as Rbc1 for <u>R</u>NA <u>b</u>inding to <u>C</u>HAP domain – <u>1</u>. Regulatory RNAs are often described as being 50-500 nucleotide long and to inhibit or activate mRNA translation, or mRNA stability through RNA base pairing to selected regions of the target mRNA (Brantl & Brückner, 2014). However, Rbc1 is much longer (> 1000) and has potential for forming an RNA duplex with the full-length *cxcA* mRNA (Fig. 3A). With Northern blotting, the Rbc1-specific probe detected transcripts of different sizes that all disappeared in the JE2 strain with a deletion of the *cxcA/rbc1* locus (Fig. 3B) confirming that all bands represent Rbc1 transcripts. In the JE2 WT, expression of *rbc1* was highest in late exponential growth (OD_600_ = 1-2) with levels declining as cells entered the stationary phase (OD ∼ 4) (Fig. 3BC). To investigate the role of Rbc1 without disrupting the *cxcA* gene, we generated a mutant lacking the three predicted promoters upstream of *rbc1* (=*rbc1(P*)*; Fig. 3A) and verified that all Rbc1 transcripts disappeared in the constructed mutant (Fig. 3B). We next measured the amount of *cxcA* transcript in the JE2 WT, and in cells lacking Rbc1 or ClpXP activity by Northern blotting (Fig. 3BC). In cells lacking either the Rbc1 transcript or the ClpXP protease, the *cxcA* mRNA levels did not change significantly (see representative Northern blot Fig. 3B, and statistics performed on three biological replicates in Fig. 3C).

**Figure 3.**
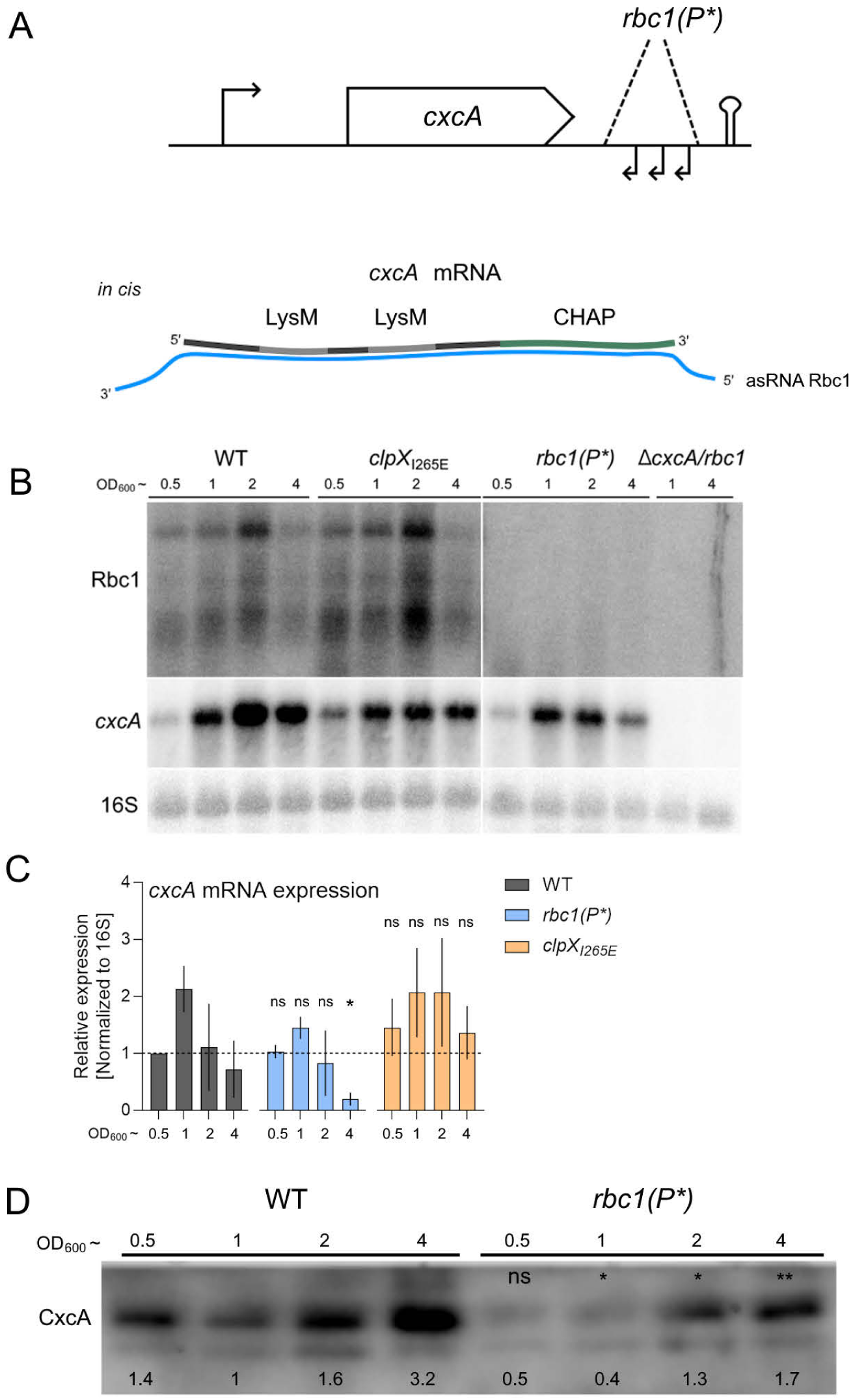
CxcA is positively controlled by the cis-encoded asRNA Rbc1. **(A)** Schematic representation showing the asRNA Rbc1 encoded from the complementary strand of the cxcA gene. Arrows indicate the three predicted promoters deleted to knock down Rbc1 transcription (this strain is designated “rbc1(P*)”. Schematic representation showing the predicted binding of the cxcA mRNA and the fully complementary Rbc1 asRNA. **(B)** Northern blot analysis performed with RNA extracted from WT or cells lacking ClpXP (clpX_I265E_), Rbc1 (rb1(P*), or CxcA and Rbc1 (ΔcxcA/rbc1) grown in TSB at 37°C until the indicated OD_600_ values. 15 µg RNA was loaded, and probed with radioactive oligos specific for Rbc1, cxcA, or 16S. **(C)** Summarized relative expression of cxcA mRNA from 3 biological replicate Northern blots, normalized to their corresponding 16S signal and compared to WT at OD_600_ ∼ 0.5. A one-tailed Welch’s t-test of unequal variance was performed for each OD_600_’s (e.g. comparing WT OD_600_ ∼ 1 to rbc1(P*) OD_600_ ∼ 1) and asterisks were given to indicate P-values: P-value > 0.05 = ns, P-value < 0.05 = *, P-value < 0.01 **. **(D)** Western blot showing CxcA levels in cell wall protein extracts for WT and cells not expressing Rbc1 at OD_600_ values 0.5, 1, 2, and 4. For statistics, three biological replicate Western blots were used and a one-tailed Welch’s t-test of unequal variance was performed for each OD_600_ category.

To investigate how knock-down of Rbc1 transcription impacts CxcA protein levels, we raised antibodies against CxcA in mice (see Methods). Interestingly, immunoblotting revealed that inactivation of Rbc1 expression is associated with a 2-fold reduction in CxcA protein levels at all OD values tested (P < 0.05; Fig 3D), with the exception of OD_600_ = 0.5. Taken together, these results suggest that CxcA levels are positively controlled by Rbc1 and negatively by ClpXP, and that regulation occurs post-transcriptionally.

### Rbc1 has potential for binding mRNAs encoding autolysins with the conserved CHAP domain *in trans*

Ten out of eighteen predicted *S. aureus* peptidoglycan hydrolases contain the highly conserved CHAP domain and the online RNA-RNA interaction tool, IntaRNA2.0 predicted that Rbc1 is capable of binding to the CHAP encoding region of mRNAs encoding the cell wall hydrolases SsaA, EssH, SAUSA300_2503, SAUSA300_0739 and Sle1 – see Table 1 and Fig. 4A (Mann *et al*., 2017; Wang *et al*., 2022). For reference, the CHAP domain part of the *cxcA* mRNA, which perfectly pairs to Rbc1, has a hybridization energy of -224.5 [kJ/mol] at 37°C, with a 100% coverage. Using IntaRNA2.0 we considered interactions with a hybridization energy of less than -25 [kJ/mol] to be possible candidate RNA-RNA interactions.

**Figure 4.**
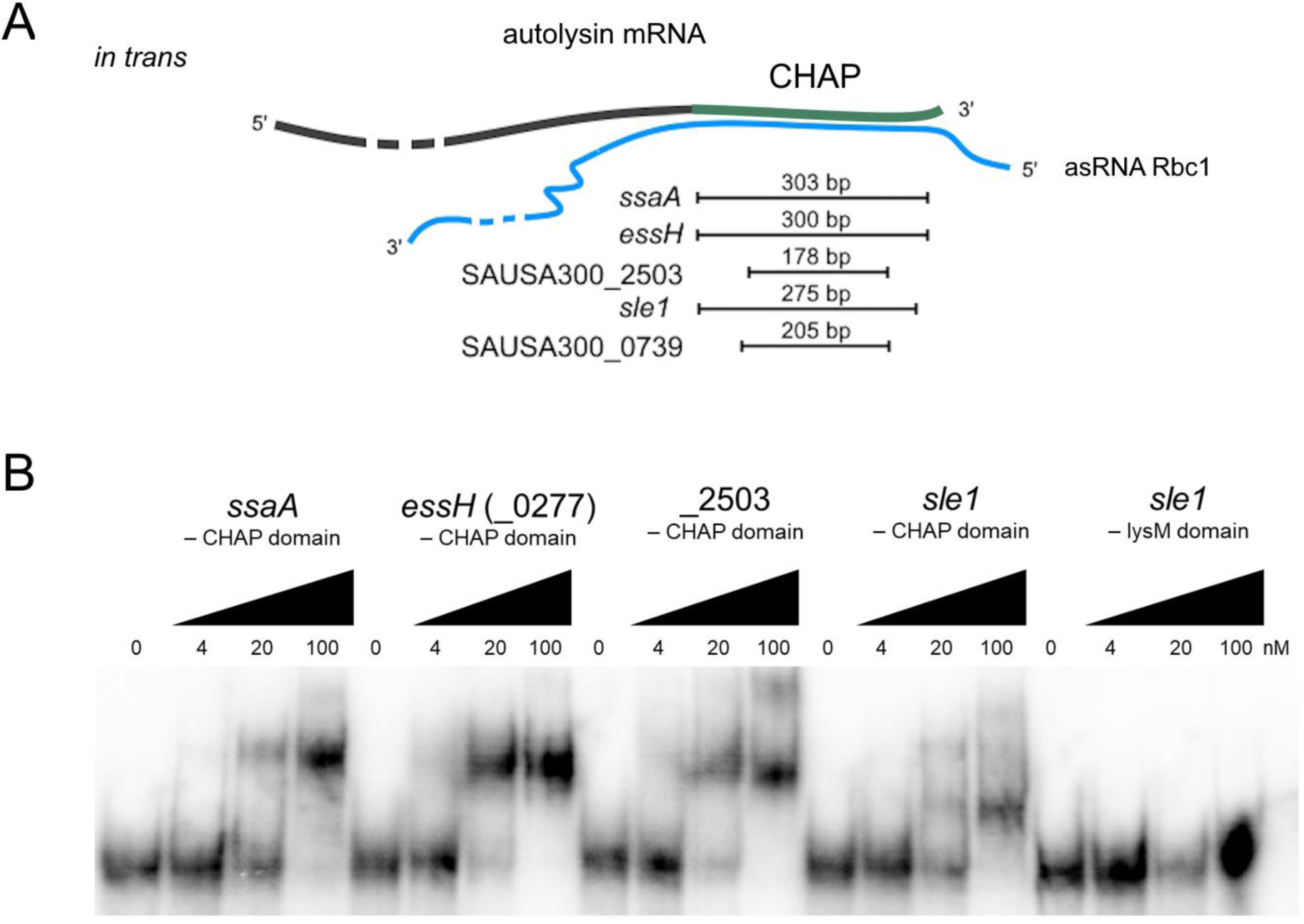
Rbc1 binds to RNAs encoding the conserved CHAP domain of cell wall hydrolases encoded in trans. **(A)** Conceptual illustration showing the predicted base pairing between Rbc1 (blue) and the CHAP encoding region of mRNAs encoding cell wall hydrolases. **(B)** Binding was verified experimentally by EMSA using 4 nm ^32^P labeled Rbc1 transcript incubated for 1 hour at 37°C with increasing concentrations of unlabeled in vitro transcribed RNA probes (200-250 nt) encoding the CHAP domains of SsaA, EssH, SAUSA300_2503 and Sle1 or, as a negative control, the LysM domain of Sle1. An upshift is indicative of interaction.

**Table 1.** Top 5 in trans-interaction partners of Rbc1 (from highest to lowest rank) predicted by IntaRNA 2.0. The hybridization energies [kJ/mol] at 37°C, calculated between Rbc1 and the full transcripts of the top 5 predicted targets of Rbc1 using IntaRNA 2.0. **\***Base pairing percentage is calculated as the ratio between the predicted number of paired nucleotides at 37°C and the section of which an interaction can be predicted – see Methods for details.

| Annotation / locus tag | Hybridization energy at 37°C [kJ/mol] | Stretch of base pairing at 37°C in Percentage* |
| --- | --- | --- |
| *ssaA* / SAUSA300_2249 | -81.1 | 230/303 – 75.9 % |
| *essH* / SAUSA300_0277 | -77.7 | 209/300 – 69.7 % |
| SAUSA300_2503 | -70.8 | 135/178 – 75.8 % |
| *sle1* / SAUSA300_0438 | -63.5 | 185/275 – 67.3 % |
| SAUSA300_0739 | -40.3 | 131/205 – 63.5 % |

The predicted interaction between Rbc1 and synthetic RNAs representing the CHAP domain encoding regions of *ssaA* (257 nt), *essH* (265 nt), SAUSA300_2503, (233 nt) or *sle1* (206 nt) mRNAs was tested *in vitro* by electrophoretic mobility shift assays (EMSA) (Lillebæk & Kallipolitis, 2024). Interestingly, the electrophoretic mobility of radiolabeled Rbc1 was clearly upshifted upon adding 20 nM or 100 nM of any of the CHAP encoding RNA fragments, while adding the same concentration of a synthetic RNA encoding the LysM domain of Sle1 (225 nt) did not cause an upshift (Fig. 4B). These results demonstrate that Rbc1 holds potential for more broadly binding to mRNAs encoding autolysins with CHAP domains, raising the possibility that Rbc1 may work *in trans* to control expression of this class of cell wall hydrolases.

### Rbc1 and ClpXP mediate temperature-dependent increase of Sle1 and CxcA protein abundance

To further explore the idea that Rbc1 is capable of controlling expression of hydrolases with a conserved CHAP domain *in trans*, we chose to focus our follow-up studies on the functionally characterized cell division hydrolase, Sle1, that has a similar modular organization as CxcA with the main difference being that Sle1 contains one additional LysM domain (Wang *et al*., 2022). The similarity between CxcA and Sle1 was supported by Western blotting showing that our CxcA raised antibody recognizes both CxcA and Sle1 as verified by the appearance of two bands corresponding in size to CxcA (28.1 kDa) and Sle1 (35.8 kDa) that disappeared in the respective mutants (Fig. 5A). RNA-RNA interactions are well known to be strongly temperature-dependent (Abduljalil, 2018; Loh *et al*., 2018) and we therefore examined the impact of Rbc1 on CxcA and Sle1 protein levels at different temperatures. Interestingly, immunoblotting revealed that abundance of the two hydrolases increases strongly with decreasing temperatures, with CxcA and Sle1 levels being 15 and 30 folds higher, respectively, in WT cells grown at 30°C compared to 42°C (Fig. 5A). Intriguingly, cells with a knock down of Rbc1 transcription harbored constitutive low levels of both Sle1 and CxcA at all temperatures demonstrating that Rbc1 is essential for the observed increase of Sle1 and CxcA at 30°C (Fig. 5A). At 30°C, Sle1 and CxcA levels were approximately 5 and 3 folds lower in the JE2 *rbc1(P*)* mutant (P < 0.01 and P < 0.001, respectively) while inactivation of Rbc1 had little effect on the protein levels of the two hydrolases at 37°C or 42°C. At the higher temperatures Sle1/CxcA levels seem to be mainly regulated negatively by ClpXP and JE2 cells expressing the *clpX_I265E_* variant, displayed a dramatic 50-100 fold upregulation of the two proteins at 42°C (Fig. 5A). A double mutant lacking both Rbc1 and ClpXP displayed constitutive high levels of both Sle1 and CxcA at all temperatures, showing that the positive effect of Rbc1 is dispensable in cells with inactivated ClpXP (Fig. 5A).

**Figure 5.**
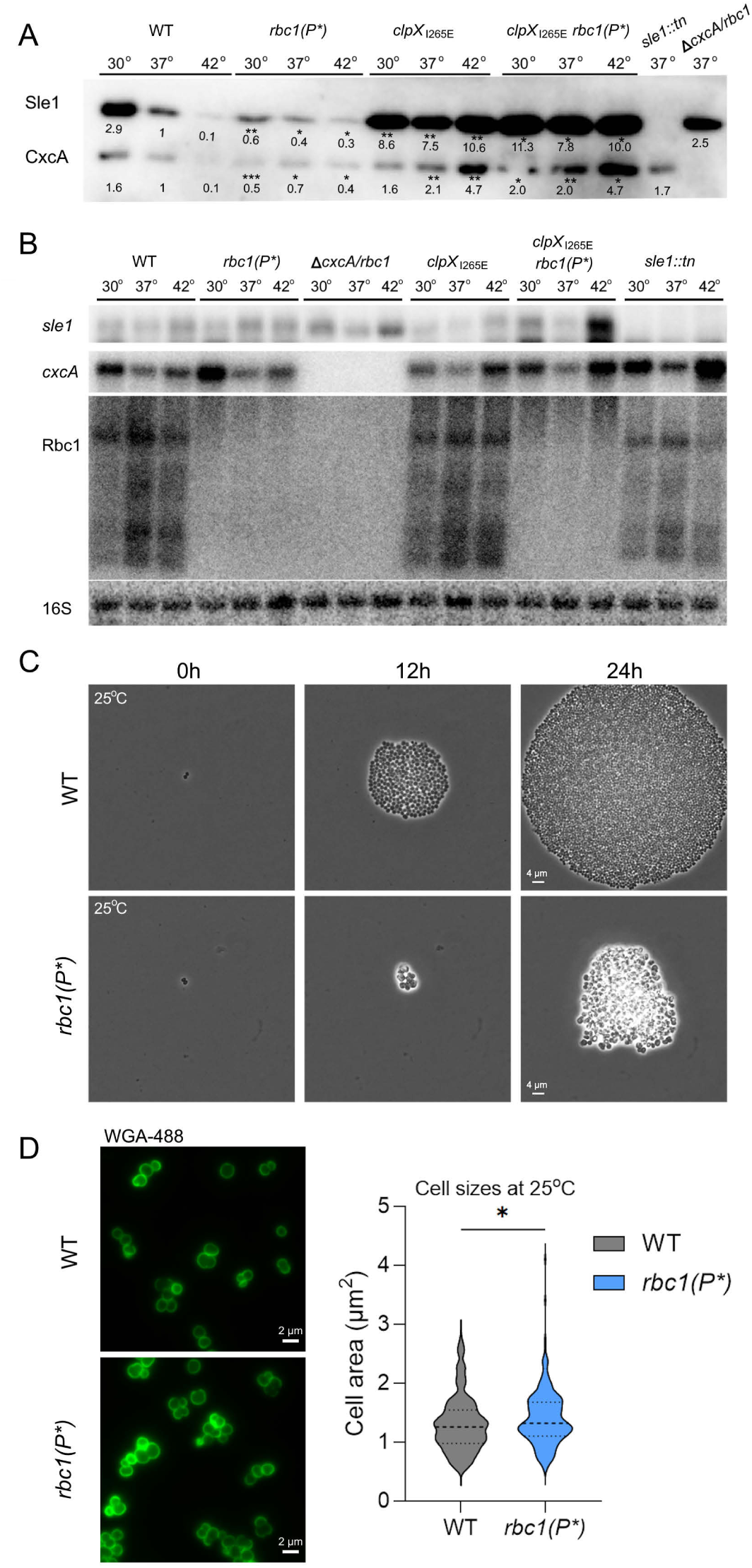
Rbc1 mediates an increase in protein levels of Sle1 and CxcA in response to decreasing temperature while ClpXP is the main regulator at 42°C. **(A)** Antibodies raised against CxcA were used in Western blotting with total protein extracts isolated from JE2 WT, JE2 rbc1(P*,) JE2 clpX_I265E_, and JE2 clpX_I265E_ rbc1(P*) cells, grown to an OD_600_ ∼ 1 at the indicated temperatures. Mutants lacking Sle1 or CxcA were included as negative controls. Band intensities were quantified using ImageLab (BioRad) and statistics were performed on the values obtained from 3 biological replicate blots, using Welch’s t-test of unequal variances and comparing values to the value for the WT sample grown at the same temperature (i.e. rbc1(P*) at 30°C compared to WT at 30°C). P-values are indicated by asterisks where * < 0.05, ** <0.01, *** < 0.001 **(B)** Northern blot analysis for the sle1, cxcA, and Rbc1 transcripts in the same strains grown under the same conditions as in (A). 16S was probed for normalization. **(C)** Simultaneous time-lapse microscopy at 25°C for the WT and rbc1(P*) strain. Cells were grown on TSB + 1.2 % agarose pads and imaged every 5 minutes, here 0, 12 and 24 hours are depicted. **(D)** WT and rbc1(P*) grown at 25°C for 4 hours before staining with WGA-488 (green) and imaged with wide-field fluorescence microscopy. Cell sizes of WT (n = 390) and rbc1(P*) (n= 348) were determined using the MicrobeJ (v. 5.13p) plugin for Fiji. Statistics were performed using a two-tailed Welch’s t-test of unequal variances, p-value = 0.0151.

To investigate if the Rbc1 mediated increase of Sle1 and CxcA occurs transcriptionally or post-transcriptionally, Northern blotting was performed under the same conditions (Fig. 5B). Most importantly this experiment showed that *sle1* and *cxcA* mRNA levels are not affected by the inactivation of *rbc1* expression at any of the tested temperatures, indicating that Rbc1 stimulates Sle1 and CxcA levels at 30°C at the post-transcriptional level. Northern blot analysis, however, revealed that while *sle1* mRNA levels did not change with temperature, the *cxcA* mRNA levels were higher at 30°C than at 37°C or 42°C, (Fig. 5B). Finally, we examined the stability of *sle1* and *cxcA* mRNA at 30°C and 37°C in JE2 WT cells with and without the Rbc1 transcript (Fig. S2). This experiment supported that Rbc1 does not impact the levels or stability of *sle1* and *cxcA* mRNA but indicated that an Rbc1-independent mechanism increased the stability of *cxcA* mRNA at 30°C, and may contribute to the upregulation of *cxcA* mRNA steady state levels at low temperature (Fig. S2).

Given the importance of Rbc1 for increasing Sle1 and CxcA protein levels at lower temperatures, we next used time-lapse microscopy to compare growth of single cells of the JE2 WT and the JE2 *rbc1(P*)* mutant at 25°C. Interestingly, the images revealed that the mutant formed colonies of reduced size and that knock down of Rbc1 transcription was associated with an aberrant cell morphology and increased clumping (Fig. 5C). To study morphology of the mutant in further detail, cells were also imaged with widefield epifluorescence microscopy after staining with the cell wall dye WGA-488 (green) that cannot penetrate the cell envelope and therefore only stains the cell wall that is exposed to the exterior. The images confirmed that the mutant cells tended to clump and appeared more irregular and heterogeneous in size, which was confirmed by estimating the cell size (Fig. 5D). This analysis also revealed that the JE2 *rbc1(P*)* mutant cells (mean = 1.397 µm^2^) are slightly larger than WT *S. aureus* cells (mean = 1.305 µm^2^, P-value = 0.0151) at this temperature (Fig. 5D). As *S. aureus* cells elongate in phase 3 (cells with closed septum), an increase in cell size is indicative of delayed autolytic splitting (Monteiro *et al*., 2015; Thalsø-Madsen *et al*., 2019).

### *S. aureus* daughter cell separation is impeded when the temperature decreases

Why would *S. aureus* need to control expression of autolysins involved in daughter cell separation according to temperature? To address this question, we used super resolution microscopy to image *S. aureus* cells grown at either 25°C or 37°C and analyzed the cell cycle by assigning cells to different phases according to the state of septal ingrowth as described above. Interestingly, SR-SIM images revealed that *S. aureus* cells grown at 25°C are significantly more frequently arranged in tetrads (Fig. 6A and 6B), a defect previously shown to arise from delayed splitting of daughter cells (Veiga *et al*., 2023). Consistent with this notion, transmission and scanning electron microscopy images of cells growing at 25°C show daughter cell pairs that have completed a second round of division in one or both of the daughter cells despite still being attached at the old septum (Fig. 6C and 6D). Tetrads are not observed at 37°C (Fig. 6A) which is consistent with the common paradigm describing that *S. aureus* daughter cells with a closed septal cross wall proceed to daughter cell splitting before initiating a new septum (Fig. 6A) (Monteiro *et al*., 2015; Zhou *et al*., 2015). At 37°C daughter cell separation occurs by ultrafast popping when the outer septal wall is degraded (Monteiro *et al*., 2015; Zhou *et al*., 2015). Of note, inward degradation of the old septal wall initiating from the periphery is frequently observed in TEM-images of tetrads from 25°C cultures indicating that resolution of the outer wall does not result in fast popping (see arrows in Fig. 6C, summarized in Fig. 6A). Taken together, these results show that splitting of daughter cells is delayed at the lower temperature, which in turn could explain why cells need to increase Sle1 and CxcA enzyme levels.

**Figure 6.**
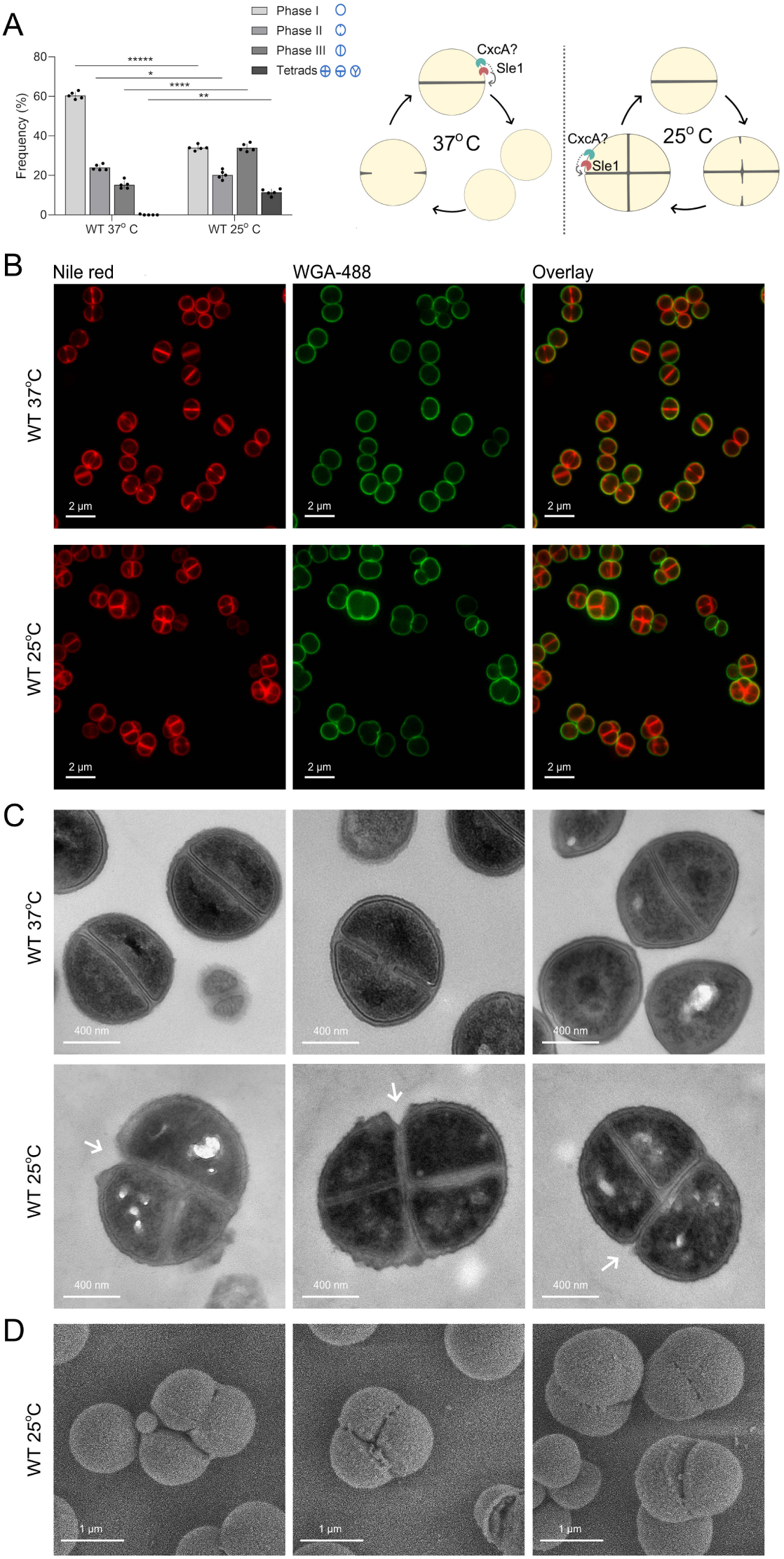
S. aureus daughter cell splitting is delayed at lower temperatures. S. aureus JE2 cells were grown at 37°C or 25°C C before imaging with SR-SIM, TEM, or SEM. **(A)** Nile Red stained cells in SR-SIM images were assigned to different phases in the cell cycle: phase I (no septum synthesis), phase II (septum synthesis started but not complete), phase III (closed septum), or tetrads (fully divided non-separated daughter cells that have completed a second round of septum formation). Cell cycle models, at either temperature, is depicted to the right. Statistics were performed using a one-tailed t-test of unequal variances (Welch’s t-test). **(B)** Representative SR-SIM images of JE2 WT grown at 37°C or 25°C. Prior to imaging cells were stained with Nile Red (membrane, red) and WGA (cell wall, green)., **(C)** TEM or **(D)** SEM images of fully divided non-separated daughter cells that have completed a second round of septum formation in one or two daughter cells in cultures of WT cells grown at 25°C.

## Discussion

In this study, we characterize the cell wall hydrolase, CxcA and show that it contributes to *S. aureus* daughter cell splitting. The hitherto uncharacterized CxcA hydrolase has an N-terminal Sec-type signal peptide and was previously shown to be among the only 8% of exoproteins that are universally expressed in all dominant clinical *S. aureus* lineages (Smith *et al*., 2016). Transcription of the *cxcA* gene is known to be under dual positive control by the GraSR and the WalKR two-component systems that are both hotspots for genetic changes associated with increasing resistance to vancomycin and daptomycin (reviewed in (Fait *et al*., 2024; Vestergaard *et al*., 2019). Transcriptional regulation might, however, not be sufficiently fast to control CxcA activity on the timeframe of a cell cycle, or in response to environmental cues, as revealed here by the discovery of two mechanisms allowing for posttranscriptional regulation of CxcA levels: i) proteolytic downregulation by the ClpXP protease, and ii) an activating mechanism involving the divergently transcribed asRNA, Rbc1. Strikingly, Rbc1 is capable of binding to the CHAP encoding region of at least four more transcripts encoding cell wall hydrolases raising the possibility that Rbc1 broadly coordinates a class of enzymes containing the same catalytic domain. In support of this hypothesis, we demonstrate that Rbc1 promotes production of the trans-encoded cell wall hydrolase Sle1, a key player in *S. aureus* daughter cell splitting (Kajimura *et al*., 2005; Thalsø-Madsen *et al*., 2019). In agreement with RNA-RNA interactions being strongly temperature dependent, Rbc1-activation of CxcA and Sle1 production is most pronounced at 30°C while vanishing at 37°C and disappearing at 42°C. In contrast, cellular levels of Sle1 and CxcA seem to be determined mainly by ClpXP-mediated proteolysis at temperatures exceeding 37°C. ATP-dependent proteolysis of functional proteins is associated with energy consumption for both synthesis and degradation and, so far, the only other protein known to be controlled by regulated proteolysis in *S. aureus* is the transcriptional stress regulator Spx, indicating that this costly type of regulation is reserved for proteins that need rapid and irreversible inactivation (Feng *et al*., 2013; Nielsen *et al*., 2024).

CxcA and Sle1 share the same modular structure with the N-terminal Sec-type signal sequence followed by, respectively, 2 or 3 peptidoglycan-binding LysM domains, and the catalytic CHAP domain. While LysM domains are ubiquitously conserved across Gram-negative and Gram-positive bacteria, the catalytic CHAP domain seems to be particularly abundant in certain Gram-positives such as staphylococci and streptococci (Mitchell *et al*., 2021). Accordingly, a protein BLAST search in the database of the National Center for Biotechnology Information (NCBI) and UniProt showed that cell wall hydrolases with full-length homology to CxcA or Sle1 (1-4 LysM domains followed by a CHAP domain) are almost exclusively found in the *Staphylococcus* genus (Fig. 7). To the best of our knowledge CxcA homologues have not been functionally characterized in other staphylococci, however, one study reported that purified *S. epidermidis* CxcA is capable of lysing *S. epidermidis* when added from the outside (Vermassen *et al*., 2019b). Staphylococcal *cxcA-*like genes, seem to always localize downstream and in opposite orientation of genes encoding the Gra-regulatory system, which respond to cell envelope stress (Fig. 7; (Falord *et al*., 2011). Interestingly, the PromoterHunter online tool predicts that Rbc1-like RNAs are transcribed from the anti-sense strand of the *cxcA* gene in *S. epidermidis, S. carnosus, S. haemolyticus,* and *S. lugdunensis*, (Fig. 7; Table 2). Therefore, the role of Rbc1 in controlling protein levels of CxcA-like cell wall hydrolases may be broadly conserved across staphylococcal species that as common colonizers of the skin and mucous membranes of animals and humans are adapted to temperatures that are lower than the core body temperature of 37°C (Bastock *et al*., 2021; Costa *et al*., 2024; Keck *et al*., 2000).

**Figure 7.**
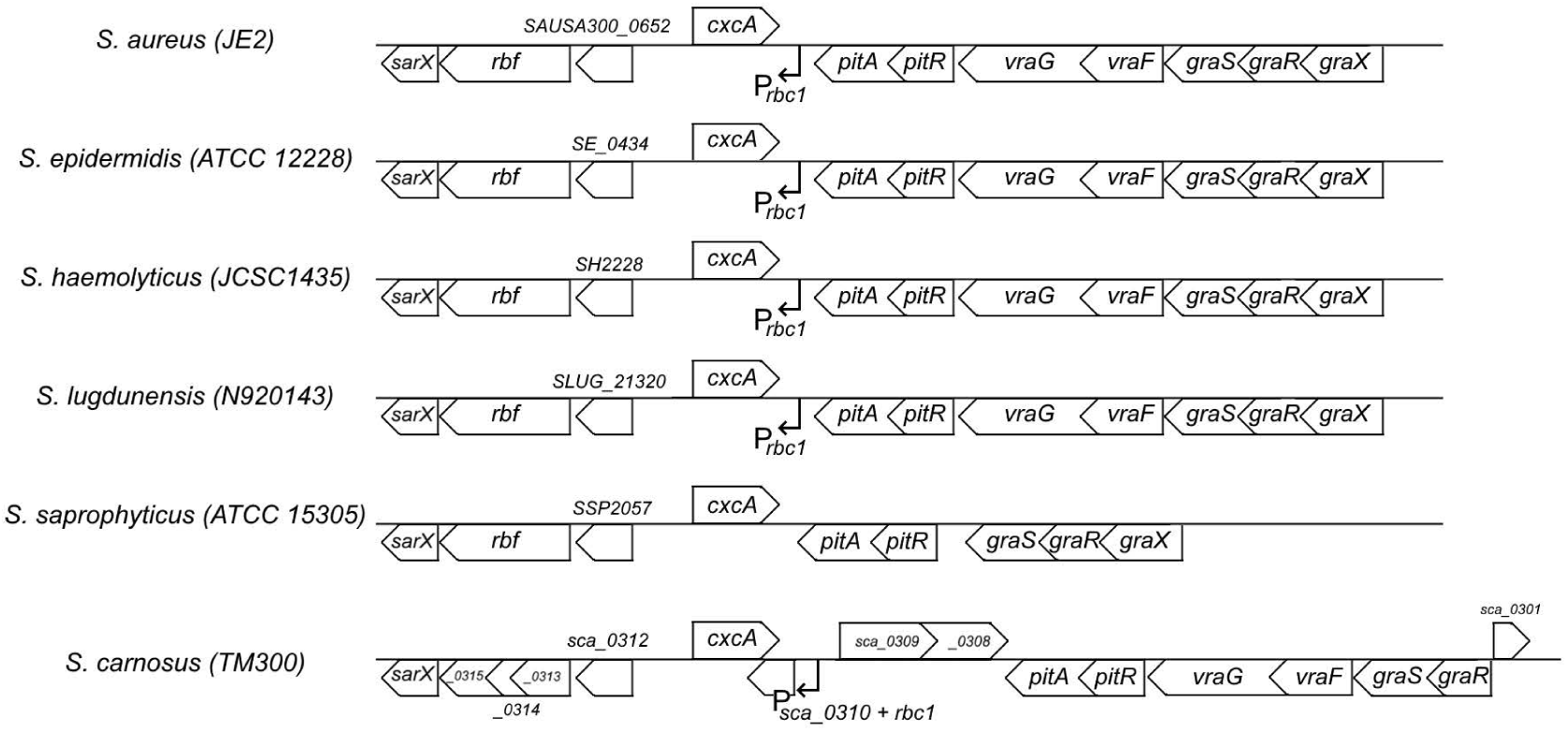
CxcA and Rbc1 homologues are encoded from a conserved genetic locus in Staphylococcal species. CxcA homologues and the adjacent gene organization illustrated for S. aureus (JE2), S. epidermidis (ATCC 12228), S. haemolyticus (JCSC1435), S. lugdunensis (N920143), S. saprophyticus (ATCC 15305), S. carnosus (TM300). The solid lined arrowed boxes indicate genes predicted by the presence of an ORF and their orientation, while the predicted rbc1 promoter is indicated by an arrow. Genes without a known name are given a locus-tag name, or simply the number of the gene, without the strain prefix, for spacing purposes.

**Table 2.** Start codons of cxcA and sle1 homologs are usually alternative codons of either UUG or GUG. Homologues of sle1 and cxcA were found using BLAST search engine. Here the start codon of the cxcA and sle1 are indicated. An rbc1 promoter could be predicted for all strains (here -35 and -10 sequences are listed), except for S. saprophyticus, using promoter matrix values for B. subtilis promoter composition (Klucar et al., 2010; Myers et al., 2021). Lastly, the analyzed genome accession number is indicated.

| Strain | Start codon of CxcA (homologs) | Start codon of Sle1 (homologs) | Predicted -35 sequence of Rbc1 | Predicted -10 sequence of Rbc1 | Genome accession no. |
| --- | --- | --- | --- | --- | --- |
| *S. aureus* (JE2) | UUG | GUG | TTGATT | TATATT | CP020619.1 |
| *S. epidermidis* | UUG | GUG | TTGTCA | TATTTT | CP035288.1 |
| *S. lugdunensis* | GUG | GUG | TTGAGT | TAATTT | CP014022.1 |
| *S. carnosus* | UUG | UUG | TTGGCA | CATAAT | NZ_CP065711.1 |
| *S. haemolyticus* | UUG | AUG | TTTAAC | TATACT | CP013911.1 |
| *S. saprophyticus* | UUG | UUG | None | None | NZ_LR134089.1 |

How does Rbc1 increase CxcA and Sle1 abundance at low temperatures? Most described RNA thermosensors localize in the 5′-untranslated region (5′-UTR) of their target genes where they control translation initiation by sequestering the ribosome binding site (RBS) in RNA secondary structures that melt at higher temperatures (Eichner *et al*., 2022). Strikingly, *in silico* prediction of RNA-based thermosensors in *S. aureus* previously identified the 5′-UTR of the *cxcA* gene (SAUSA300_0651) as a top candidate for being a conventional thermosensor raising the possibility that *cxcA* mRNA folds into a translationally inactive form at lower temperature (Hussein *et al*., 2019). A scenario could therefore involve Rbc1, which by an unknown mechanism, transforms *cxcA* mRNA into a translation competent form at 30°C. This again raises the question of why *S. aureus* would invest resources in such complex regulations. Generally, the activity of PG hydrolases needs to be strictly coordinated with peptidoglycan synthesis at all growth conditions to ensure cell wall integrity. The relative rate of cell wall synthesis and degradation likely varies with temperature, and as shown here, *S. aureus* daughter cell separation is delayed relative to septum synthesis at 25°C. Therefore, *S. aureus* cells need a mechanism to upregulate cell division hydrolases without compromising integrity of the cell wall, and we speculate that the *cxcA* 5′-UTR hairpin prevents unsynchronized translation of the *cxcA* mRNA. In bacteria, translation of most mRNAs initiates while transcription is still ongoing, however, transcription and translation may be uncoupled in genes that require very tight regulation to avoid cell death. Classical examples are the widespread type I toxin– antitoxin (TA) systems consisting of a toxic protein and a noncoding asRNA acting as an antitoxin by preventing translation of the toxin through direct base pairing between the asRNA and the toxin mRNA (Bonabal & Darfeuille, 2024). Of relevance here, the toxin mRNA is initially folded into a translation-incompetent structure that secure that translation is not initiated prior to binding to the antitoxin RNA (Bonabal & Darfeuille, 2024). Moreover, toxins are often translated from non-canonical, less efficient start codons that function to delay translation initiation and secure time for base pairing between the toxin mRNA and the asRNA (Bonabal & Darfeuille, 2024). Intriguingly, we noted that Sle1 and CxcA across staphylococcal species are encoded from the non-canonical start codons UUG and GUG, respectively (Table 2). In *Escherichia coli*, the UUG start codon is used in less than 1 % of genes and is estimated to have only 10% of the activity of the conventional AUG start codon (Hecht *et al*., 2017). With inspiration from the TA-systems described above, we speculate that the use of poor start codon generally delays translation initiation of *cxcA* and *sle1* mRNAs, which could create time for binding of Rbc1. Specifically, the proposed RNA-based thermosensor in the 5′-UTR of the *cxcA* gene in combination with the UUG start codon could prevent untimely translation of upregulated *cxcA* mRNA at the lower temperature. To prevent cell lysis, the activity of cell division hydrolases must be tightly coordinated with septum synthesis, and we hypothesize that binding of Rbc1 to *cxcA* and *sle1* mRNAs plays a role in coordinating translation of these hydrolases with septum synthesis in space and time. Our data does, however, not support that Rbc1 is required for translation per se as knock down of *rbc1* increased Sle1 and CxcA levels in cells devoid of ClpXP (Fig. 6A). Therefore, an alternative model is that Rbc1 promotes increased abundance of Sle1 and CxcA via mitigating proteolysis (Fig. 8). Along the same line, a recent study revealed that the divisomal DNA translocase protein FtsK protects Sle1 from being degraded by ClpXP and that Sle1 was completely degraded in cells lacking FtsK (Veiga *et al*., 2023). Interestingly, we noted that also CxcA is absent from the published proteome of the *ftsK* mutant (Veiga *et al*., 2023). Together, these finding indicate that a common mechanism involving the divisomal FtsK protein is protecting Sle1 and CxcA from being degraded by the ClpXP protease. Presumably, such a mechanism allows *S. aureus* to coordinate the activity of cell division hydrolases with DNA replication and progression of septum synthesis (Fig. 8).

**Figure 8.**
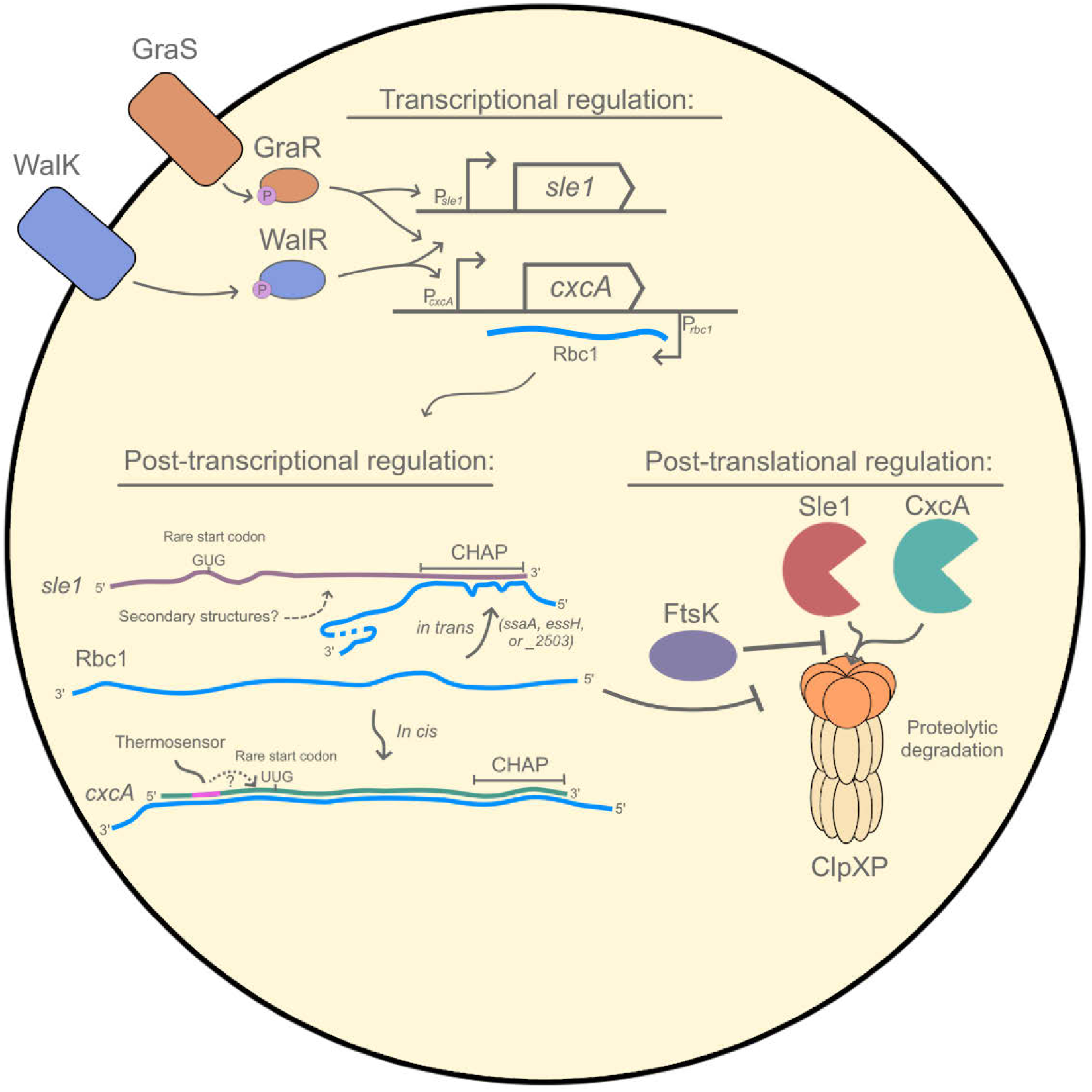
Overview and proposed model for sle1 and cxcA regulation on the transcriptional, post-transcriptional and post-translational level. **Transcriptional regulation:** the cxcA and sle1 genes are both regulated by the two-component systems WalKR and GraSR, where the response regulators WalR or GraR, upon phosphorylation bind to the promoter (P) of sle1 or cxcA to activate transcription of the genes. The regulatory RNA, Rbc1 is transcribed from the opposite strand of the cxcA gene. **Post-transcriptional regulation:** The Rbc1 asRNA binds in cis to the cxcA mRNA and in trans to the CHAP domain region of the sle1 mRNA (or ssaA, essH or SAUSA300_2503). To allow time for Rbc1 binding, transcription and translation are uncoupled through mechanism that delay translation initiation, including the folding of the 5’ end of the cxcA mRNA into a temperature-dependent secondary structure that covers the Shine-Dalgarno sequence (Hussein et al., 2019). Likewise the sle1 mRNA can be computationally predicted using the PASIFIC online tool (Millman et al., 2017). Additionally, translation is delayed by the use of rare, inefficient start codons UUG in cxcA mRNA, and GUG in the sle1 transcript. **Post-translational regulation:** Sle1 and CxcA are substrates of the ClpXP protease (Feng et al., 2013). FtsK protects and CxcA from ClpXP degradation during the cell cycle (Veiga et al., 2023). Moreover, Rbc1 has a role in protecting CxcA and Sle1 from ClpXP proteolysis, particularly at low temperatures. See main text for details.

Due to their lethal potential, the activities of bacterial cell wall hydrolases must be precisely regulated through tight transcriptional, translational, and post-translational controls. In conclusion, we here uncover novel mechanisms allowing for post-transcriptional control of cell division hydrolases in the bacterial pathogen *S. aureus* (Fig. 8). At temperatures ≥37°C, cellular levels of CxcA and Sle1 are primarily controlled by ClpXP proteolysis. At lower temperatures, daughter cell separation is delayed, and abundance of Sle1 and CxcA is upregulated post-transcriptionally by Rbc1 transcribed from the antisense strain of the *cxcA* gene. Rbc1 binds to the CHAP domain encoding region of *sle1* mRNA as well as other mRNAs encoding the catalytic CHAP domain and further studies will explore if Rbc1 more broadly coordinates protein levels of cell wall hydrolases with the conserved catalytic CHAP domain. Interestingly, a non-coding asRNA (S989) is transcribed from the complementary strand of the *ssaA* gene encoding another conserved *S. aureus* cell wall hydrolase with the CHAP domain (Mäder *et al*., 2016). Moreover, the 3′UTR of the *vigR* gene was recently shown to post-transcriptionally promote expression of the cell wall lytic transglycosylase *isaA* through direct mRNA–mRNA base-pairing. Therefore, RNA-RNA interactions may broadly be used to control expression of cell wall hydrolases in *S. aureus* and related bacteria. (Mediati *et al.,* 2022).

## Materials and Methods

### Cultivation and Transformation

Strains of *Staphylococcus aureus* JE2 MRSA WT and mutant derivatives were routinely grown on Tryptic Soy Agar (TSA) plates or in Tryptic Soy Broth (TSB) liquid medium at 37°C degrees or indicated otherwise. Mutagenesis was performed using allelic exchange or transduction. For transformation, cells were cloned using the pIMAY-Z vector system, allowing for selection with chloramphenicol (20 µg/mL), and subsequent blue-white screen of mutants, as described in (Monk & Stinear, 2021). The *cxcA* knockout mutant was created by in-frame deletion of *cxcA*, retaining the first 10 and last 12 amino acids of CxcA. The *rbc1(P*)* inactivation mutant lacks 344 nucleotides of the promoter-region of *rbc1*, located adjacent to *cxcA*, while retaining the predicted terminator sequence of *cxcA*. Primers with overhangs containing KpnI (Primer ID with an A prefix) or NotI (Primer ID with a D prefix) restriction sites were used for insertion into the pIMAY-Z vector. Primer sequences used for creating the mutants are listed in Table 3. Promoters were predicted by using the online tool, PromoterHunter, with standard parameters, finding 3 promoters for *rbc1* (Klucar *et al*., 2010). For double-mutants creation, the *clpX_I265E_* allele with an erythromycin selection cassette, was transduced into *ΔcxcA/rbc1* and *rbc1(P*)* background strains using bacteriophage Φ11, selecting on agar plates with erythromycin concentrations of 7.5 µg/mL.

**Table 3.**
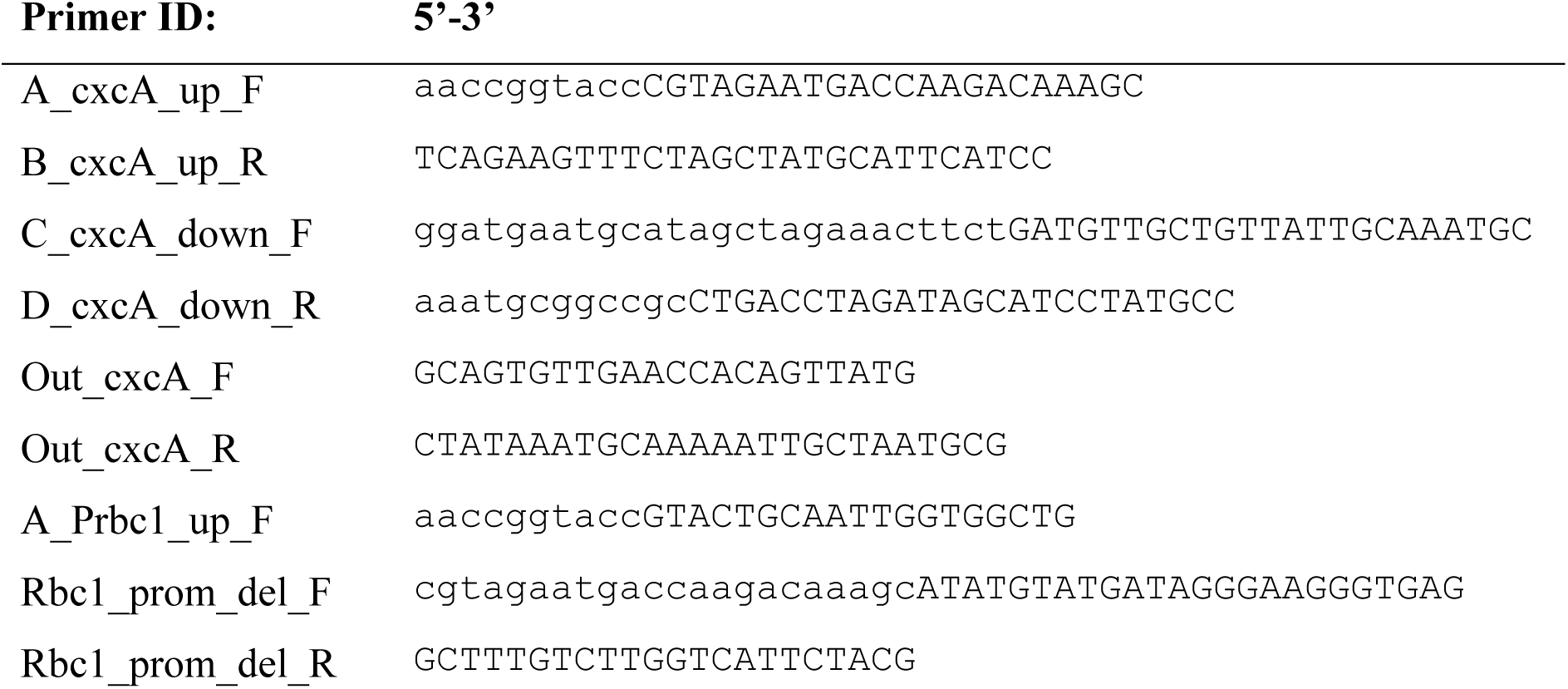
Primer list used for creating the ΔcxcA and rbc1(P*) mutants. Upper-case letters indicate annealing regions, while lower-case letters indicate non-annealing overhangs.

### Cultivation and sample harvest for proteomic analysis

A ClpXP deficient strain was constructed in the NCTC8325-4 background to identify ClpXP substrates. Here the NCTC8325-4 Δ*clpX* (Frees *et al*., 2003) was complemented using the pSK9067 plasmid (Brzoska & Firth, 2013) expressing either a functional wild type *clpX* sequence or a *clpX_I265E_* sequence. The resulting ClpX variant proteins are C-terminally His_6_-tagged. ClpX interaction with ClpP is diminished for the ClpX_I265E_ protein by mutation of the IGF-loop motif resulting in ClpXP protease deficiency but no ClpX chaperone deficiency in *S. aureus* cells carrying the *clpX_I265E_* variant (Stahlhut *et al*., 2017). The NCTC 8325-4 Δ*clpX* pSK9067_clpX and pSK9067_clpX_I265E_ were cultivated aerobically at 37°C in pMEM (1x MEM for SILAC; 1x NEAA; 4 mM L-glutamine; 10 mM HEPES; 2 mM alanine, valine, leucine, isoleucine, aspartate, glutamate, serine, threonine, cysteine, proline, phenylalanine, histidine and tryptophan; sterile filtrated; pH 7.4). For the SILAC approach, the medium was supplemented with 0.576 mM light arginine and 0.556 mM light lysine for the mutant variant carrying strain whereas the wild type variant carrying strain was cultivated in medium supplemented with 0.0576 mM heavy arginine and 0.556 mM lysine (^13^C_6_ Arg-HCl and ^13^C_6_ Lys-2HCl). The medium was prepared as described in (Pförtner *et al*., 2013). The medium was additionally supplemented with 10 µg/ml erythromycin for plasmid selection and 800 µM isopropyl-β-D-thiogalactopyranoside (IPTG) for induction of *clpX* variant expression.. Four biological replicates of *S. aureus* cells expressing the *clpX* variants were aerobically precultured in medium supplemented with 0.01% (m/v) yeast extract at 37 °C and exponentially growing samples were used for inoculation of the main culture to an optical density at 600 nm (OD_600_) of 0.05. At the exponential growth phase (OD_600_ = ca. 0.4) and at early-stationary phase (OD_600_ = ca. 1.8), 20 OD units were harvested by cooled in liquid nitrogen and subsequent centrifugation at 17,105 x g, 3 min, 4 °C. Cell pellets were washed with 1 ml of cold 10 mM HEPES with 50 mM NaCl pH 8.0 and stored at -80 °C after snap freezing in liquid nitrogen.

### Sample preparation for mass spectrometry measurements

The bacterial cell pellets were thawed on ice and wild type variant and mutant variant carrying cells were joined per biological replicate and growth phase in 500 µl of 4°C 10 mM HEPES with 50 mM NaCl pH 8.0. Cells were mechanically disrupted using 200 µl glass beads (Sigma) in the FastPrep-24 (4x 45 sec 600 rpm, 300 sec cooling on ice in between). Then, 5 U/ml of benzonase (Pierce^TM^ Universal Nuclease, Thermo Fisher Scientific) was added to the cell lysates and the lysates were incubated for 10 min. Cell debris was removed by centrifugation at 16,200 x g for 1h at 4°C. Protein concentration of the protein lysates was determined by the Micro BCA^TM^ Protein-Assay-Kit (Thermo Fisher Scientific). Tryptic digestion of proteins and subsequent peptide purification were performed as described in detail in Blankenburg *et al*., 2019, with minor changes as described in Ganske *et al*., 2024. 4 µg protein mixture per (mutant and wild type mixed) sample were incubated with 80 µg of hydrophilic (GE Healthcare, Little Chalfont, UK) and hydrophobic (Thermo Fisher Scientific, MA, USA) carboxylate-modified magnetic SeraMag Speed Beads in 70% (v/v) acetonitrile (ACN). During incubation beads were shaken in a ThermoMixer (1,400 rpm, RT, 18 min). Beads were washed two times with 70% (v/v) ethanol, one time with 100% (v/v) ACN and afterwards dried. Then ACN-free beads were incubated with 160 ng trypsin (Promega Corporation, Wisconsin, USA) in 20 mM ammonium-bicarbonate buffer (37 °C, 16 h) for tryptic digestion of the sample proteins. The digestion was stopped by addition of ACN (final 95% (v/v)) and shaking in a ThermoMixer (18 min, 1400 rpm, RT). Beads were washed with 100% (v/v) ACN and then peptides were eluted from the beads with 2% (w/v) dimethyl sulfoxide and ultra-sonication for 5 min. Finally, the samples were resuspended in buffer A (4% (v/v) ACN, 0.2% (v/v) acetic acid) and stored at -20 °C.For nano LC-MS/MS analysis, tryptic peptides were separated on an Ultimate 3000 nano-LC system (Thermo Fisher Scientific, MA, USA) and subsequently analyzed on an Orbitrap Exploris^TM^ 480 mass spectrometer (Thermo Fisher Scientific, MA, USA) in data-independent acquisition (DIA) mode. Further details of the MS/MS-analysis are summarized in Table S2-AB and S3.

### Data analysis of mass spectrometry data

DIA-MS data was searched using the Spectronaut software (version 18.7.240506.55695; Biognosys AG, Switzerland) against a dedicated spectral library specifically constructed for this project including DIA-MS measurements of several *clpX* allele variant expressing strains comprising of 20,774 precursors and an *S. aureu*s NCTC 8325 protein data base (fasta file retrieved from *Aureo*Wiki in December 2017 comprising of 2852 staphylococcal proteins and five contaminant proteins) (Fuchs *et al*., 2018). For spectral library construction, the Spectronaut software (version 14.8.201029.47784) using the Pulsar search engine was exploited. Detailed parameters for the search are summarized in Table S3.

Subsequently, the Spectronaut-intern median-normalized ion-ratio data was analyzed using R (v4.1.2). Packages exploited for the analysis are listed in Table S4. Protein identifiers were associated with the meta-information and gene symbols as retrieved from *Aureo*Wiki (Fuchs *et al*., 2018) in June 2024. Peptide ratios were calculated as median of ion ratios per peptide and sample and likewise, protein ratios were calculated as median of peptide ratios per protein and sample. Protein ratios were global median-normalized. Statistics on differentially abundant proteins were calculated using the limmaDE approach (Phipson *et al*., 2016; Ritchie *et al*., 2015). P-values were adjusted for multiple testing (q-value) according to Benjamini and Hochberg (Benjamini & Hochberg, 2018). Proteins, which were detected with at least two peptides and an absolute fold change of equal or more than 2 and a q-value equal or less than 0.05 were considered as significantly different in abundance.

The mass spectrometry proteomics data have been deposited to the ProteomeXchange Consortium *via* the PRIDE partner repository with the dataset identifier PXD053967 (Perez-Riverol *et al*., 2024).

### Reviewer access details

Log in to the PRIDE website using the following details:

**Project accession:** PXD053967

**Token:** FnjtrOZ4TUuC

Alternatively, reviewer can access the dataset by logging in to the PRIDE website using the following account details:

**Password:** xELhezLShMdR

### Simultaneous Time-lapse Microscopy

For visualization of *S. aureus* colony morphology, we performed simultaneous time-lapse microscopy. Strains of WT, *clpX_I265E_*, *clpX_I265E_* Δ*cxcA/rbc1* and *rbc1(P*)* were grown in TSB media for 2 hours at 37°C, before adjusted to OD_600_ ∼ 0.05. A 2.8x1.7cm Gene Frame (Thermo Fischer Scientific), was attached to a clean microscope slide. A 4-well construction was created within the attached Gene Frame, using a clean scalpel attaching fragments of another Gene Frame. TSB media supplemented with 1.2 % agarose, was melted at 84°C for 30-60 minutes in a heating block. 1 mL melted TSB-agarose mixture was added to the Gene Frame, avoiding air bubbles, and a clean microscope slide was applied on top of the TSB agarose, to level the TSB agarose mix. The mixture was left to solidify at room temperature for 2-4 minutes, while light pressure was applied on top of the Gene Frame. The slide was then carefully removed. Using a clean scalpel, the agarose mix in the septal areas was removed. The plastic of the Gene Frame, was then removed, exposing the sticky side of the Gene Frame. A microliter of diluted *S. aureus* culture (OD_600_ ∼ 0.05) was added to each individual well and was slightly air dried. A 2.4x3.2 cm coverslip was applied on top aligning with the Gene Frame and sealed by running the bottom of a pen along the edges. Phase-contrast images were acquired using an Olympus IX83 inverted motorized microscope. The temperature of the microscope was set to 25°C within the cellSens incubator module. Using the cellSens software, 3 positions for each strain were saved, and a program was created whereby the microscope localized each position every 5 minutes. At the pre-saved positions, two autofocus steps were applied before phase-contrast imaging. The first autofocus step was set for a wider range (40 µm in Z-direction) with coarse stepping of 2 µm and fine stepping with 1 µm. The second autofocus step was set at small range (15 µm in Z-direction), with a 1 µm coarse stepping and 0,5 µm fine stepping. The ROI of the autofocus steps were modified in close proximity to the micro colony. Images were acquired every 5 minutes for 48 hours, using a 100x 1.4 N.A oil-immersion objective and a sCMOS Photometrics Prime camera. The shutter was closed between each iteration to avoid heating of the sample. Images were later analyzed in ImageJ and a video produced herein.

### Super Resolution – Structured Illumination Microscopy (SR-SIM)

Cells were grown till mid-log phase at indicated temperatures in TSB liquid medium. Cells were stained with Nile Red (10 µg/mL), WGA-488 (2µg/mL) and HADA (250 µM) and spotted on PBS 1.2 % agarose pads within a Gene Frame (Thermo Fischer Scientific). SR-SIM was performed with an Elyra PS.1 microscope (Zeiss) using a Plan-Apochromat 63x/1.4 oil DIC M27 objective and a Pco.edge 5.5 camera. Images were acquired with five grid rotations and reconstructed using ZEN software (black edition, 2012, version 8.1.0.484) based on a structured illumination algorithm, using synthetic, channel specific optical transfer functions and noise filter settings ranging from -6 to -8. SR-SIM was performed at the Core Facility for Integrated Microscopy, Faculty of Health and Medical Sciences, University of Copenhagen.

### Immunization of mice to CxcA

For raising the CxcA/Sle1 antibody, *E. coli* strain BL21 was transformed with an IPTG inducible vector, pET28a, carrying a synthetic construct of 6xHis-tag fused to CxcA at both the N- and C-terminus. For protein production, cells were grown in 2L Lysogeny Broth (LB) with 50 µg/mL kanamycin for selection. At mid-log phase, cultures were shifted to 16°C and 500 µM IPTG was added, before overnight incubation. Cells were subsequently pelleted and lysed using French Press, and the protein was purified using His-GraviTrap (Cytiva) columns. CxcA was purified according to manufacturer’s instructions, but eluted using a gradient of imidazole 20, 40, 50 and finally 200 µM imidazole. The purity of the eluted fractions was assessed using SDS-PAGE and Coomassie staining. The pure protein fraction was repeatedly buffer exchanged using a 10 kDa spin-filter (Amicon) in appropriate buffer for immunization. Purified protein was lyophilized and sent for mouse immunization at Evaxion Biotech, Denmark.

### Immunoblotting of cell wall associated protein

Cultures of 50 mL were used for cell wall protein isolation. Here, cultures were pre-grown overnight at 37°C and diluted to OD_600_ ∼ 0.05 and grown until indicated OD_600_ values at indicated temperatures. Cultures were pelleted by centrifugation at 7.000 x g for 5 minutes at 4°C and washed once in cold 0.9% NaCl. Pellets were resuspended in 10 mL 4% SDS and incubated at 25°C for 45 minutes with shaking. Cells were then pelleted and supernatant was transferred to 1:1 volume cold 96% ethanol and precipitated overnight at 4°C. Precipitated cell wall proteins were pelleted by centrifugation at 4°C, 10.000 x g for 30 minutes. Supernatant was discarded and protein pellet was dried in a fume hood. Protein pellets were normalized to the cultures OD_600_ value and resuspended in 0.5xOD_600_ mL 300 mM NaCl, 20 mM HEPES pH 7.4 and 1 % SDS. Proteins were separated by gel electrophoresis, using a NuPage 4-12 % SDS-PAGE pre-cast gel, and subsequently transferred onto a 0,22 µm PVDF membrane. Membrane was blocked in 5 % Skim milk PBS-Tween for 1 hour and incubated in 1:4.000 primary mouse CxcA-antibody in 2 % BSA and PBS-Tween overnight. Membrane was washed in PBS-Tween before adding secondary HRP conjugated mouse antibody (Dako) in 2 % skim milk PBS-Tween for 1 hour. The membrane was then washed 3x with PBS-Tween, before addition of Immobilon Forte HRP substrate (Merck). Substrate was applied briefly and imaged on an Amersham ImageQuant 800 GxP.

### RNA purification and Northern blotting

Cells were grown as indicated, before 6-10 mL of culture was snap frozen with brief submersion (∼ 10 seconds) into liquid nitrogen before being pelleted in a pre-cooled centrifuge. The pellet was preserved at -70°C until RNA purification. For RNA purification, cells were resuspended in 700 µL Na-Acetate buffer (Na-Acetate pH 4.5, 1 mM EDTA, 0.5 % SDS) and transferred to a FastPrep tube with acid washed glass beads. Cells were lysed by 3x 40s 6.0 m/s shaking in a Homogeniser, FastPrep-24 (MP Biomedicals) and placed on ice between each iteration. RNA was purified using a hot phenol-chloroform approach. The cell lysate was centrifuged (3 min, 5.000 x g) and supernatant was transferred to a tube with 100 µL chloroform and 500 µL acidic phenol (pH 4.5). The mixture was heat-treated at 65°C for 10 min with shaking in a heating block (1100 rpm), inverting the tubes every 2-3 minutes followed by brief vortexing. The mixture was centrifuged and the aqueous phase was transferred to a tube containing 500 µL chloroform and vortexed for 20s. The mixture was centrifuged again and the aqueous phase transferred to another tube with 500 µL chloroform and vortexed. The sample was centrifuged and the aqueous phase was transferred to 1:3 volume of 96% cold ethanol with 1:10 volume of 3M Na-Acetate (pH 4.5) and left precipitating at -20°C overnight. RNA was pelleted by centrifugation at 20.000 x g at 4°C for 45 min and washed once with 96% ethanol and twice with 70% ethanol. The pellet was dried in a fume hood and resuspended in nuclease free H_2_O. RNA integrity was assessed on a TAE agarose gel with EtBr, and RNA-concentrations were measured using NanoDrop. 15 µg of RNA was mixed with TriTrack 6x DNA loading dye (Thermo Fischer Scientific) and formamide (w/v of 25 %) and heated at 80°C for 5 minutes before loading onto a 10 mM NaPi and 1 % agarose gel. Gels were run at 75 V for 2.5 hours, and transferred to a ZetaProbe (BioRad) membrane by capillary blotting in 20xSSC buffer overnight. RNA was crosslinked and briefly washed with autoclaved milliQ-H_2_O and dried in a fume hood. Membranes were probed with radioactive ∼ 40 nt oligos, specific for either Rbc1, *cxcA*, *sle1* or 16S. Here 0.3 µM oligo was routinely labeled with ^32^P using PNK kinase. For membrane probing, 10 mL hybridization buffer with 2-4 µL radioactive probe was incubated overnight at 42°C in a rotation incubator. After overnight hybridization, the membrane was washed twice, in 2xSSC citrate buffer, and 0.5xSSC citrate buffer for 10 minutes each. The membrane was dried in a fume hood and placed onto a phosphor screen overnight. Images were acquired by scanning the phosphor screen with a Typhoon FLA 9500.

### IntaRNA2.0 for screening potential RNA-RNA binding partners

For RNA-RNA interaction prediction, we utilized the online tool, IntaRNA2.0 (Mann *et al*., 2017). As IntaRNA2.0 only accepts 750 bp, we screened the first 750 nts of Rbc1 pairing to all replicons of the NCBI RefSeq: NC_007795 (*Staphylococcus aureus* NCTC 8325). Here we used the 1-300 nt upstream and downstream of the replicon stop (or start) codon. The remaining parameters were left with standard settings.

Upon initial screening of potential targets in NCTC 8325, we manually inserted the corresponding RNA-sequences from the JE2 strain, allowing for longer interaction prediction exceeding 300 nts from the stop (or start) codon.

### Electrophoretic Mobility Shift Assay (EMSA)

EMSA was performed as described in (Lillebæk & Kallipolitis, 2024). T7 promoters were attached to PCR products of Rbc1*, cxcA, sle1, ssaA, essH* and SAUSA300_2503. PCR products of 200-250 bp were created with High-Fidelity Phusion polymerase (New England Biolabs) and DNA was purified via GFX DNA purification kit (Thermo Fischer Scientific) according to manufacturer’s instructions. Single-stranded RNA was synthesized using the T7 MEGAScript kit according to manufacturer’s instructions (Thermo Fischer Scientific) and purified by separation with a 6 % denaturing poly-acrylamide gel. RNA bands were cut from the gel and incubated in 2M ammonium acetate pH 5.5, and purified with subsequent addition of phenol pH 4.5, proceeding with a phenol-chloroform RNA purification. The Rbc1 single stranded RNA transcript was phosphatase treated with Shrimp Alkaline Phosphatase (Sigma Aldrich) and radioactively labeled with ^32^P using Polynucleotide Kinase (New England Biolabs) according to manufacturer’s instructions. Single-stranded Rbc1 RNA was purified using Nucleospin miRNA kit (Macherey-Nagel) according to manufacturer’s instructions. For EMSA, native 6 % poly-acrylamide gel was cast and pre-run for 30-45 minutes at 90V in a 4°C cold room. Tubes for EMSA was prepared of desired mix of labeled Rbc1, yeast tRNA, and target RNA in a KCl, HEPES buffer for a final volume of 10 µL and incubated at 37°C for 1 hour. Samples were mixed with 5 µL 50% glycerol and loaded onto the gel while running. The gel was run for 2.5 hours in a 4°C cold room. The gel was carefully separated from the glass plates, and dried in a gel drier for 1-1.5 hours. The dried gel was placed in a phosphor screen overnight and imaged using a Typhoon FLA 9500.

### Fluorescence microscopy and image analysis

*S. aureus* cultures pre-grown overnight at 37°C, were diluted to OD_600_ ∼ 0.05 and grown at 25°C for 4 hours. Cells were stained with 10 µg/mL Nile Red and 2 µg/mL WGA-488 for 5 minutes prior to washing 2x with PBS. Cells were spotted on PBS 1.2 % agarose pads inside a Gene Frame (Thermo Fischer Scientific). Cells were imaged with a Photometrics Prime sCMOS camera using a 100x 1.4 N.A phase contrast objective in an inverted Olympus IX83 microscope. Phase-contrast and fluorescent images were acquired in succession.

Images were loaded into Fiji (v. 216/1.54p) and subsequently analyzed using the MicrobeJ plugin (v. 5.13p) (Ducret *et al*., 2016). Cells were detected using the phase-contrast image with the following settings: Area: 0.5-5 µm^2^, Length: 0-5 µm, Width: 0-5 µm, Circularity: 0.7-max. Exclude on Edges, Shape descriptors, Segmentation, Straighten, Shape, and Profile were checked before loading into the manual editor. Here, cells incorrectly joined, were segmented before loading into the results tab. Here, SHAPE.area was used as the measured area and plotted in GraphPad Prism (v.10.4.1).

### Transmission electron microscopy (TEM)

*S. aureus* cultures pre-grown at 37°C, were diluted (OD_600_ ∼ 0.05) into 40 ml of TSB and grown at 25°C or 37°C until OD_600_ ∼ 0.5. 10 mL culture volume was pelleted by centrifugation at 8,000 x g, washed in PBS pH 7.4, and re-suspended in fixation solution (2.5% glutaraldehyde in 0.1 M cacodylate buffer [pH 7.4]) and left overnight at 4°C. The fixed cells were further treated with 2% osmium tetroxide, followed by 0.25% uranyl acetate for contrast enhancement. The pellets were dehydrated in increasing concentrations of ethanol, followed by pure propylene oxide, and then embedded in Epon resin. Thin sections for electron microscopy were stained with lead citrate and observed in a Philips CM100 BioTWIN transmission electron microscope fitted with an Olympus Veleta camera with a resolution of 2,048 by 2,048 pixels. Sample processing and microscopy were performed at the Core Facility for Integrated Microscopy, Faculty of Health and Medical Sciences, University of Copenhagen.

### Scanning electron microscopy (SEM)

Similarly to TEM imaging, *S. aureus* cells were grown to mid-exponential phase and collected by centrifugation, washed in PBS and fixed in 2.5% glutaraldehyde with 0.1 M cacodylate buffer, pH 7.4. The fixed cells were left overnight at 4°C. They were then carefully resuspended and sedimented onto coverslips for 1 week at 4°C. The cells were then washed three times in 0.15 M sodium phosphate buffer, pH 7.4 and samples were post-fixed in 1% OsO_4_ and 0.12 M sodium cacodylate buffer, pH 7.4 for two hours. Samples were rinsed in distilled water, before progressively dehydrated into 100% ethanol and point dried (Balzers CPD 030) with CO_2_. Samples were afterwards mounted on stubs using a sputter coated with 6 nm gold (Leica Coater ACE 200) and colloidal silver as adhesive. Samples were imaged with a FEI Quanta 3D scanning electron microscope operated at an accelerating voltage of 2 kV. Sample preparation and SEM imaging was performed at the Core Facility for Integrated Microscopy, Faculty of Health and Medical Sciences, University of Copenhagen.

