## Supplemental Figures for "Temperature-dependent regulation of bacterial cell division hydrolases by the coordinated action of a regulatory RNA and the ClpXP protease"

### Supplementary figures:

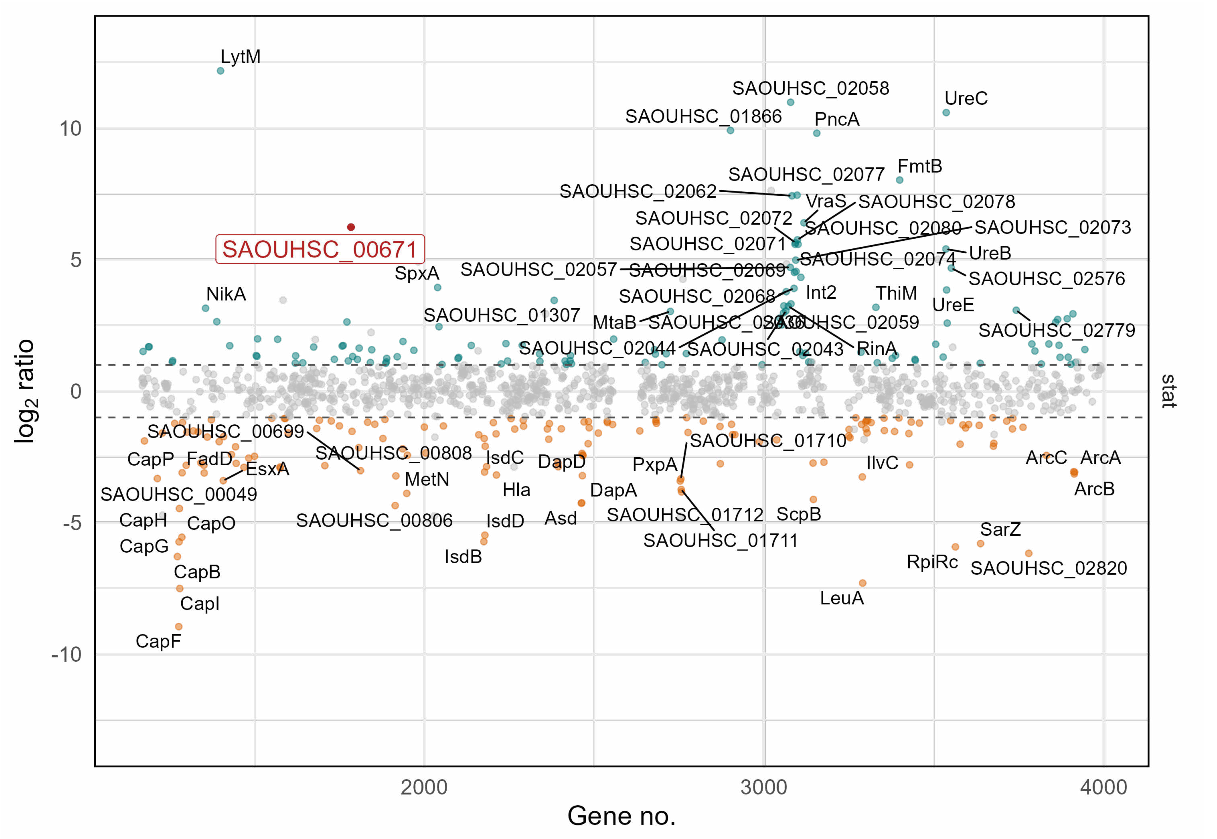


***Figure S1| SAOUHSC_00671 and Sle1 are upregulated in a clpX$_{I265E}$ mutant.*** *Proteomics of stationary phase samples of NCTC8325-4 WT and its mutant derivative clpX$_{I265E}$ show that SAOUHSC_00671 is increased in stationary phase samples. Unique proteins are plotted with a dot (grey if Benjamini-Hochberg adjusted p-value > 0.05 and/or log2 ratio is < 1 and/or > -1 and a peptide count > 2 cutoff) according to their genomic placement coordinate. Proteins, which are significantly changed in abundance (adjusted p-value < 0.05 and log2 ratio > 1 or log2 ratio < -1) are given either blue (upregulated genes) or brown (downregulated genes). Significantly changed proteins with a absolute log2 ratio > 3 are labeled. SAOUHSC_00671 is marked in red).*

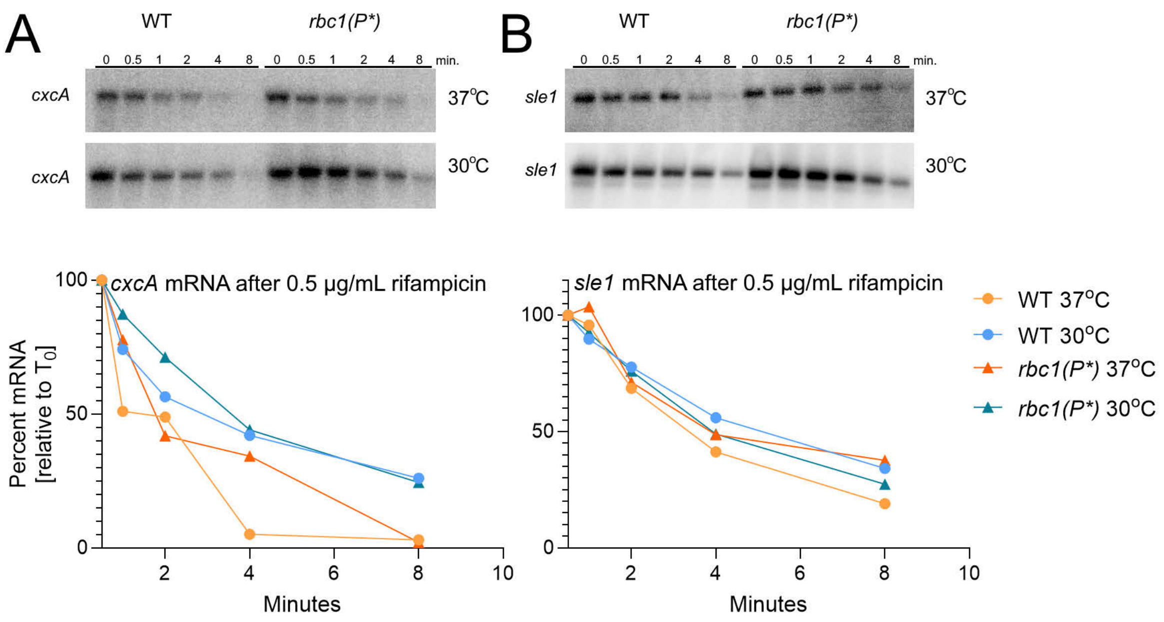


***Figure S2| cxcA and sle1 mRNA stability, is unaffected by Rbc1 inactivation. Temperature however extends the half-life of cxcA.*** *Cells were grown until OD ~ 1, before adding 0,5 mg/mL rifampicin. Samples were obtained at 0, 30 seconds, 1, 2, 4 and 8 minutes at either 37°C or 30°C. Sample images of Northern blots probed for either* ***(A)*** *cxcA or* ***(B)*** *sle1, are displayed above extinction curves. Extinction curves are displayed as percentage from* $T_0$ *versus time (minutes). Circles indicate wild-type cxcA or sle1 levels (orange for 37°C and blue for 30°C), while triangles for the rbc1(P*) mutant (red for the 37°C and green-blue for the 30°C).*
